# Topological data analysis reveals principles of chromosome structure throughout cellular differentiation

**DOI:** 10.1101/540716

**Authors:** Natalie Sauerwald, Yihang Shen, Carl Kingsford

## Abstract

Topological data analysis (TDA) is a mathematically well-founded set of methods to derive robust information about the structure and topology of data sets, and has been applied successfully in several biological contexts. Derived primarily from algebraic topology, TDA rigorously identifies persistent features in complex data, making it well-suited to better understand the key features of three-dimensional chromosome structure. Chromosome structure has a significant influence in many diverse genomic processes and has recently been shown to relate to cellular differentiation. While there exist many methods to study specific substructures of chromosomes, we are still missing a global view of all geometric features of chromosomes. By applying TDA to the study of chromosome structure through differentiation across three cell lines, we provide insight into principles of chromosome folding and looping. We identify persistent connected components and one-dimensional topological features of chromosomes, and characterize them across cell types and stages of differentiation.

**Availability:** Scripts to reproduce the results from this study can be found at https://github.com/Kingsford-Group/hictda

## 1 Introduction

The three-dimensional shape of chromosomes has significant influence in many critical cellular processes, including gene expression and regulation [7, 14, 26], replication timing [24, 2], and overall nuclear organization [10]. The wide range of processes related to chromosome structure suggests that understanding this component is crucial to a broader explanation of many genomic mechanisms, yet it remains a challenge to study this complex system.

In particular, the process of differentiation, by which a cell changes to a new cell type, is critical to all multi-cellular life, but the mechanisms behind this process remain an active field of research with recent work suggesting a role for chromosome structure in this process. Structural changes have been observed in chromosomes through lineage specification, both across several stages of human cardiogenesis [16] as well as across human embryonic stem cells (ESCs) and four human ES-cell-derived lineages [12]. Fields et al. (2017) [16] identified both global and local structural dynamics, observing transitions from repressive to active compartments around cardiac-specific genes as they are upregulated through differentiation. Dixon et al. (2015) [12] also noted structural dynamics across hierarchical scales during development, with some corresponding gene expression changes.

Chromosome structure can be measured by a number of variants of the chromosome conformation capture protocol [11], including Hi-C [21] which permits genome-wide measurements of the chromosomal architectures of a population of cells. Hi-C quantifies physical proximity by counting cross-linkage frequencies between genomic segments. Because of the dependence on cross-linking, which is likely to induce both false positive and false negative contacts, the heterogeneity within cell populations, and the large scale and complexity of the system, Hi-C can be very challenging to analyze. Many methods have focused on identifying local structures [17], however it has proven challenging to study large-scale structures across the entire genome.

A class of techniques called “Topological Data Analysis” (TDA) has gained prominence recently as a generalized, mathematically grounded set of methods for identifying and analyzing the topological and geometric structures underlying data. Emerging from work in applied topology and computational geometry, TDA aims to infer information about the robust structures of complex data sets [9]. These methods have already been applied to various biological contexts [3], including in studies of gene expression at the single cell level [27], viral reassortment [8], horizontal evolution [4], cancer genomics [23, 1], and other complex diseases [20, 18]. Similar methods have also been used in tools to enable large-scale biological database searching [31]. The two main methods of TDA are Mapper, a dimensionality reduction framework and visualization method, and persistent homology, an algorithm for extracting geometric and topological structures which describe the underlying data.

Given its rigorous mathematical foundation and ability to identify important topological structures, TDA is very well-suited to the analysis of Hi-C data. Emmett et al. (2015) [15] first applied these methods to human Hi-C data, though computational limitations at the time restricted this analysis to only one chromosome at 1Mb resolution. More recently, TDA was used to analyze the similarities between single-cell Hi-C maps [6]. Carriere and Rabadan (2018) [6] first computed pairwise distances between all single-cell Hi-C contact matrices, and applied TDA to the distance matrix between single cells rather than applying TDA directly to the Hi-C data, analyzing the results with Mapper. In this paper, we use persistent homology to identify geometric structures in human chromosomes directly from Hi-C data, and study how they change throughout lineage specification and differentiation.

This work presents the first use of TDA to study the chromosome structures of all 22 human autosomal chromosomes, providing insight into the structural changes involved in cellular differentiation. We identify hundreds of thousands of zero- and one-dimensional features of chromosome structure across 14 cell types, and describe the patterns underlying geometric structures of Hi-C data. We characterize the one-dimensional hole structures identified by TDA, noting that many of the patterns we observe can be explained by the linearity of the chromosome. Additionally, we compare the topologies of 14 cell types representing various stages of differentiation and various cell lines, and note that the topological similarity is largely dictated by cell line rather than differentiation stage.

## 2 Methods

### 2.1 Overview of TDA

TDA is based on the premise that data points are sampled from an unknown continuous geometric structure that can be described by topological properties preserved under continuous deformations of the space. These properties can include the number and size of connected components, loops or holes the structure contains. TDA approximates a continuous geometry by building a *simplicial complex*, or a network of edges and triangles, from the nodes of the given data points. More complex simplices of *n* dimensions can be generated for high-dimensional data, but for the purposes of Hi-C, which is only three dimensional, our simplicial complexes are made up only of simplices of dimension at most 2, i.e., nodes, edges, and triangles. A Vietoris-Rips (VR) complexis then generated, which is a set of simplices produced by adding edges between all nodes with distance less than a given *α*, and a triangle between all sets of three nodes for which each pair is no more than *α* apart. Together these components describe a structure built from the data, from which the topological features of the underlying space can be described and quantified through a process called *persistent homology*.

#### Definition: Vietoris-Rips complex

Given a set of points *X* in a metric space (*M, d*) and a real number *α* ≥ 0, The Vietoris-Rips complex is the set of simplices{[*x*_0_, …, *x*_*k*_]} such that *d*(*x*_*i*_, *x*_*j*_) *≤ α* for all (*i, j*), with *k* less than or equal to a given maximum dimension [9].

The analysis of simplicial complexes and their topological properties is based in homology theory, which defines the topological properties of any given dimension of a space. These properties can be represented by homology groups *H*_0_(*X*), *H*_1_(*X*), *H*_2_(*X*), …, *H*_*n*_(*X*). A homology group *H*_*k*_ represents *k*-dimensional “holes”. For example, *H*_0_ represents the connected components of the VR complex, *H*_1_ represents one-dimensional loops, and *H*_2_ represents two-dimensional voids [29].

Given a set of data points *X*, we build VR complexes for different values of parameter *α*. The basis of persistent homology is the idea that features that persist in the VR complexes across values of *α* are the key topological features of the space generated by the data. Conversely, features that only appear for few values of *α* are often filtered out or ignored as methodological artifacts. A feature from persistent homology is therefore described by an interval [*b, d*], where *b* represents the birth time of the feature, or the smallest value of *α* at which the feature is found, and *d*, called death time, the smallest value of *α* at which the feature no longer exists. These features are visualized in two ways: *persistence diagrams* and *barcode plots*. Persistence diagrams are sets of (*b, d*) points in the Euclidean half-plane above the diagonal. Barcode plots display the same information, but with each homology group shown as an interval [*b, d*]. These barcode lines are plotted at different heights, often ordered by death time *d* for visualization purposes, but the *y* values are arbitrary. For more technical details on TDA and persistent homology, see for example Carlsson (2014) [5] or Wasserman (2018) [30].

### 2.2 Applying TDA to Hi-C

The methods of TDA use a distance matrix that describes the distances between all data points. Although Hi-C data is interpreted as describing the 3D distances between chromosomal segments, the values of a Hi-C matrix are contact counts rather than distance values, where a high contact count implies a low distance. We use the following transformation to convert a normalized Hi-C matrix *M* to a distance matrix *K*:

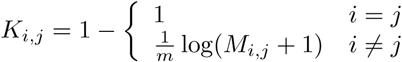

where *m* = 1.01 max_*i,j≤D*_(log(*M*_*i,j*_) + 1), and *D* is the number of rows in the contact matrix. A pseudo-count of 1 is added to all off-diagonal values in the Hi-C matrix to avoid taking a logarithm of zero, and the factor of 1.01 is included to ensure that all distances where *i* ≠ *j* are nonzero. Note that this transformation does not return a metric, as Hi-C values themselves do not satisfy metric properties. The practical impact of violating the metric assumption in TDA has not been explored, so it is unclear how this influences our results.

We use GUDHI [22], a Python library for TDA, to compute persistent homology from these distance matrices at 100kb resolution, with the maximum dimension of simplices created is 2.

### 2.3 Loop trace-back algorithm

The *H*_1_ structures identified by TDA represent “loops” in the input data, or one-dimensional holes in the simplicial complex. We will use the term loop interchangeably with *H*_1_ structure but it is important to note that these are not loops in the traditional sense of chromatin loops. The loops identified by TDA may be surrounded by non-consecutive genomic segments, unlike the continuous loops generally studied in chromatin.

TDA does not identify the data points involved in each *H*_*k*_, as each element corresponds to a homology class rather than an individual structure. Each *H*_1_ class represents the set of loops around a one-dimensional topological hole, not a specific loop structure in the data. In order to locate a representative loop for each homology class, we first define an “optimal loop” as the loop in the homology class with the shortest total distance. In the Hi-C case, loop edges are connections between two chromosome bins *i* and *j*, and their edge weights are given by *K*_*i,j*_ in the distance matrix. We therefore look for the loop which minimizes the sum of edge weights over all loops in the homology class.

In order to identify the optimal loop of each homology class *H*_1_ we use the following loop trace-back algorithm, based on identifying a shortest path through the distance matrix *K*, similar to the method used by Emmett et al. (2015) [15]. One major difference in our algorithm is that we do not perturb the Hi-C matrices, ensuring that we preserve all spatial relationships among genomic loci. Besides, we provide a proof of the correctness of our loop trace-back algorithm(S1 in the supplementary file). The formal loop trace-back algorithm is described as follows:

- Step 1: For each homology class in *H*_1_ from the persistent homology, identify the unique edge which forms at the birth time and generates the homology class. We will refer to the two nodes of this edge as *V*_1_ and *V*_2_.
- Step 2: Construct a weighted graph *G* from the distance matrix, where nodes are chromosome bins and edges are weighted by the distance *K*_*i,j*_, including only edges where *K*_*i,j*_ is less than the distance between *V*_1_ and *V*_2_.
- Step 3: Find the shortest (lowest weight) path from *V*_1_ to *V*_2_ on the graph using Dijkstra’s algorithm, and return this path plus the edge between *V*_1_ and *V*_2_.

From the proof we can see that this algorithm is based on two assumptions: First, that there is only one homology class formed by an edge *e* and second, that there are no other edges from the original distance matrix with the same length as *e*. In the Hi-C case, these are not strong assumptions: all of the homology classes generated here satisfy the first assumption and only around 2% of them do not satisfy the second one. The cases violating the second assumption have only one other edge with the same weight, indicating that this loop trace-back algorithm will perfectly identify almost all optimal loops for our data but may not be suitable for other applications.

### 2.4 Null models

In order to understand the TDA loop structures, three separate null models of distance matrices representing various properties of the Hi-C data were also analyzed and compared to the *H*_1_ structures of the original Hi-C data. The three null models are defined as follows:

- Random permutation: all Hi-C distance values are permuted randomly, preserving only the symmetry of the distance matrix and the values themselves.
- Edge permutation: the distance values along each row of the distance matrix were permuted randomly, preserving both the degree of and set of distances for each node but randomly changing their assignments. The corresponding columns were permuted in the same way to preserve symmetry.
- Linear dependence: a new distance matrix is created, in which each diagonal beyond the main diagonal preserves the same mean and standard deviation of the original data, with Gaussian noise added. The dominant pattern of the Hi-C distance matrices is a decrease in distance as the difference between the row index and column index increases. This model represents this same pattern, but does not include any additional structural features of chromosomes.

### 2.5 Bottleneck distance to compare persistence diagrams

In order to derive stability results for TDA, Carlsson (2014) [5] proposed a metric called the *bottleneck distance* that quantifies the difference between two persistence diagrams. The bottleneck distance is based on a perfect bipartite matching *g* between two persistence diagrams dgm_1_ and dgm_2_, where points in either persistence diagram can also be matched to any point along the diagonal. The formula for computing the bottleneck distance *d*_*B*_ is:

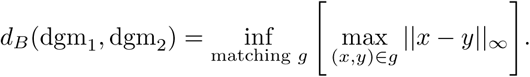

The bottleneck distance quantifies the similarity between two persistence diagrams by the maximum distance between two points in the best matching.

## 3 Results

### 3.1 Data

We analyzed Hi-C samples from 14 conditions, representing several differentiation lineages from two different studies [16, 12]. Two of these lineages represent paths through human cardiogenesis, starting with stem cells and continuing through the mesoderm (MES), cardiac progenitor (CP), and cardiac myocyte (CM) stages. One line, from RUES2 cells, begins with embryonic stem cells (ESC), and also includes a fetal heart tissue sample. The study authors also collected data from WTC11 cells, beginning with a human induced pluripotent stem cell (PSC), then collecting data at the same stages as the RUES2 cells: MES, CP and CM [16]. The third Hi-C data set represents still another differentiation starting point, using H1 ESC cells to generate four human ES-cell-derived lineages: mesendoderm (ME), mesenchymal stem (MS) cells, neural progenitor (NP) cells, and trophoblast-like (TB) cells [12]. All data is described in Table 1, including accession codes. Samples from all 14 conditions included two replicates each. All of the Hi-C data was processed from raw reads to normalized contact matrices at 100kb using the HiC-Pro pipeline [28] and iterative correction and eigenvector decomposition (ICE) normalization [19]. In order to maximize coverage, we combined all of the reads from replicates to produce one Hi-C matrix per sample.

**Table 1:**
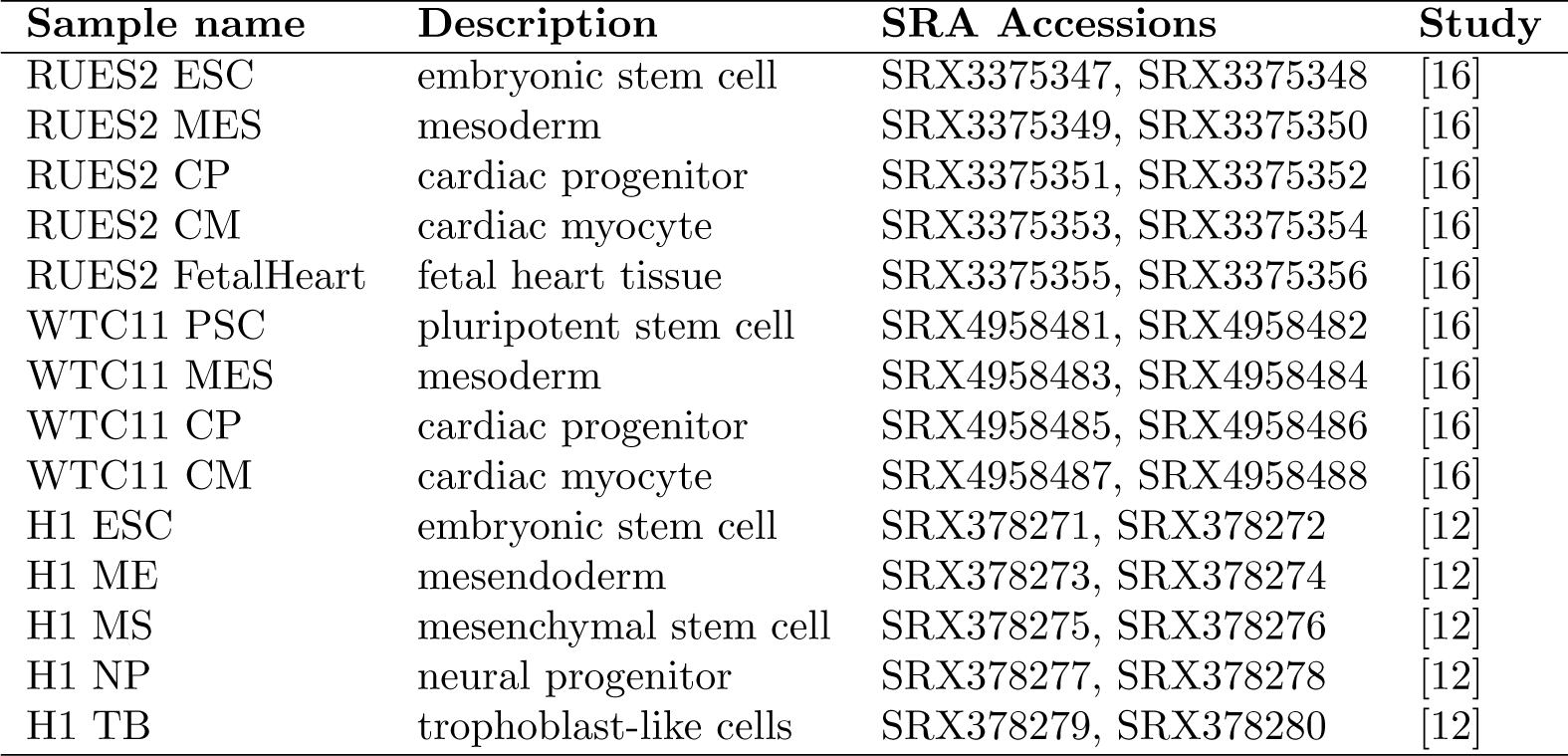
All Hi-C samples used for this study.

We generated ten of each of the null models for every RUES2 cell type and each chromosome from 13 to 22. The null models on longer chromosomes proved not to be computationally feasible, but the patterns across the chromosomes that we were able to model were remarkably consistent (see Figures S16, S17, and S18), suggesting that the additional data from all three null models on chromosomes 1 through 12 would follow similar patterns.

### 3.2 Persistent homology in Hi-C data

We visualize the persisent homology groups in the two ways described previously, persistence diagrams and barcode plots. We focus here on *H*_0_ and *H*_1_ structures and observe a very distinctive pattern in both structure classes across chromosomes and cell types in all of our Hi-C data. The majority of *H*_0_ structures dis-appear at a radius of somewhere between *α* = 0.1 and *α* = 0.2, suggesting that many new edges are formed near these values, and relatively few *H*_0_ structures persist after this. TDA therefore quickly recovers the linear structure of the chromosome. The *H*_1_ structures tend to have short lifespans (they are close to the diagonal in the persistence diagrams), and most are born at *α* ∼ .0.6 –.0.8,though there are consistently a few loops born earlier. A representative barcode plot and persistence diagram can be seen in Figure 1, and barcode plots for all samples on all autosomal chromosomes can be seen in Figures S2–S15.

**Figure 1:**
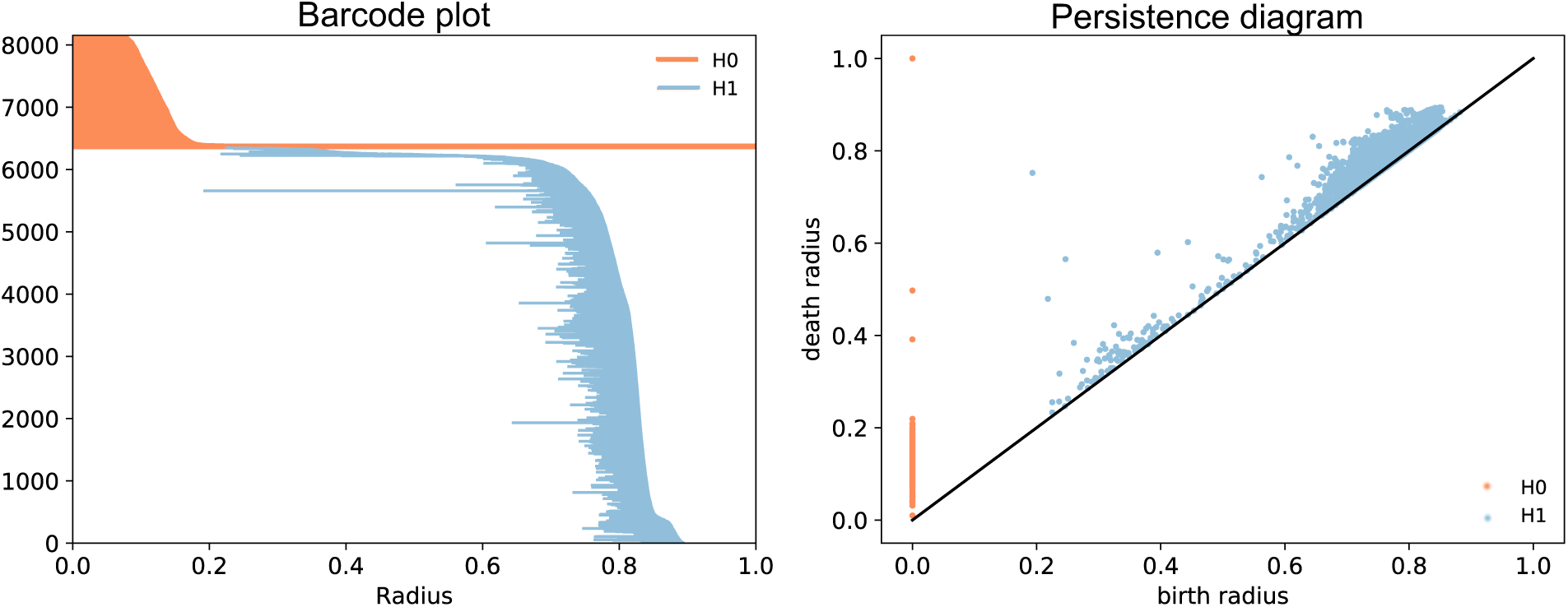
Representative example of a barcode plot and persistence diagram from Hi-C data. These figures were created from chromosome 5 of WTC11 cardiac progenitor cells, and summarize the output from the persistent homology computation. Each point or bar represents one structure, defined by the radius at which the structure can first be seen (birth radius), and the last radius before the structure no longer exists (death radius). These figures are two ways to represent the same persistent homology information.

### 3.3 Characterization of loops

Using the loop trace-back algorithm described in Methods, we locate the representative genomic bins for each *H*_1_ homology class identified and characterize their distributions across cell type, chromosomes, and differentiation stages. Recall that TDA “loops” are not traditional chromosome loops in the sense that they may be made up of non-consecutive genomic segments; across all of our data only 1868 of the 850188 total loops found are made up of consecutive genomic segments with mean length of only ∼625kb. Given that most of the loops are not made of consecutive genomic segments, we will refer to their size as the number of bins involved in the loop rather than the genomic distance between the first and last bin. These loop structures are generally very small (mean ∼446.5kb, or 4.47 bins), but this mean is largely dominated by very short-lived loops; 98.98% of all *H*_1_ structures identified by TDA have a lifespan (difference between birth and death radius) of less than 0.1, which can be seen in both the barcode plot, where the majority of blue lines are very short, and the persistence diagram, where most blue dots are very close to the diagonal (Figure 1). We therefore look at these loop size distributions as a function of their lifespan (Figure 2a), noting that while the average loop is quite short, there are many much longer loops (the longest loop involves 2492 bins, equivalent to 24.92Mb) and the distributions vary significantly between lifespan ranges. Removing the loops with lifespan less than 0.1 which are likely artifacts, the distributions of loop size do not vary much between cell types (Figure 2b), though there is some variation between chromosomes (Figure 2c). We did not observe any consistent trends in loop characteristics through differentiation across the three cell lines (Figure S1).

**Figure 2:**
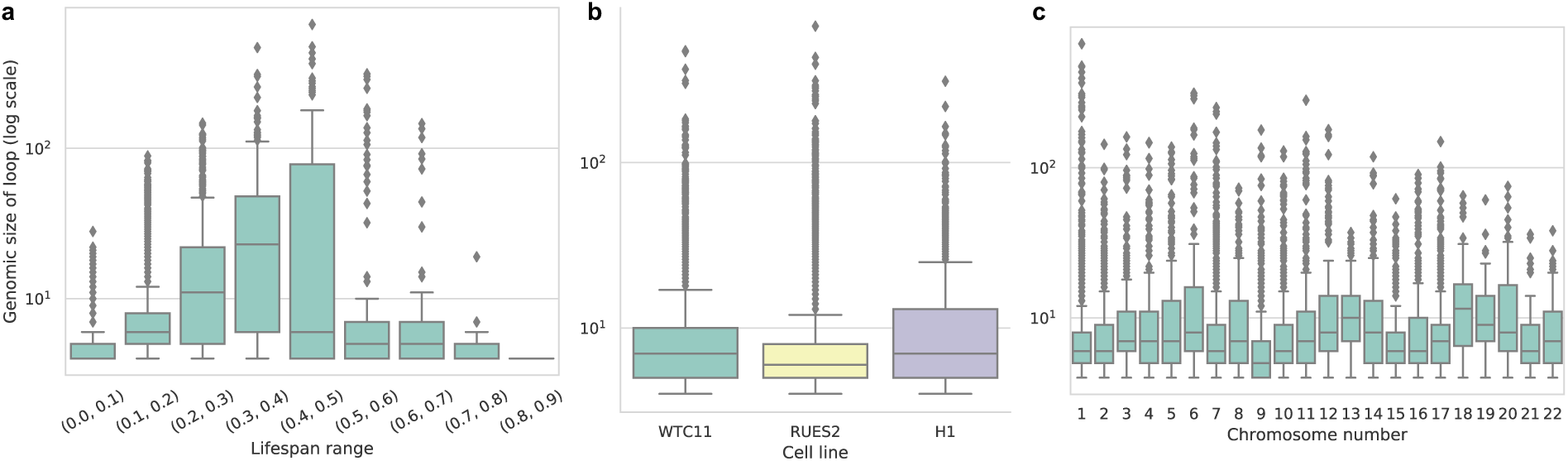
Distributions of genomic loop sizes across loop lifespans, cell types, and chromosomes. For all three figures, the log-scaled y-axis represents the total number of genomic bins returned by the loop traceback method. (a) Loop distributions separated by their lifespans in the VR complex. Larger lifespans indicate loops that are present for a large range of *α* values. (b) Loop size distributions by cell type, excluding all loops with lifespans less than 0.1, which are likely to be artifacts. (c) Loop size distributions by chromosome, again excluding all loops with lifespans less than 0.1.

### 3.4 Loop patterns are largely dictated by the linearity of chromosomes

In order to further understand the *H*_1_ structures, the patterns observed in real Hi-C data were compared to our null models. As a representative example, barcode plots of all loop structures of chromosome 14 in RUES2 MES cells can be seen in Figure 3a, along with the null models from this data.

**Figure 3:**
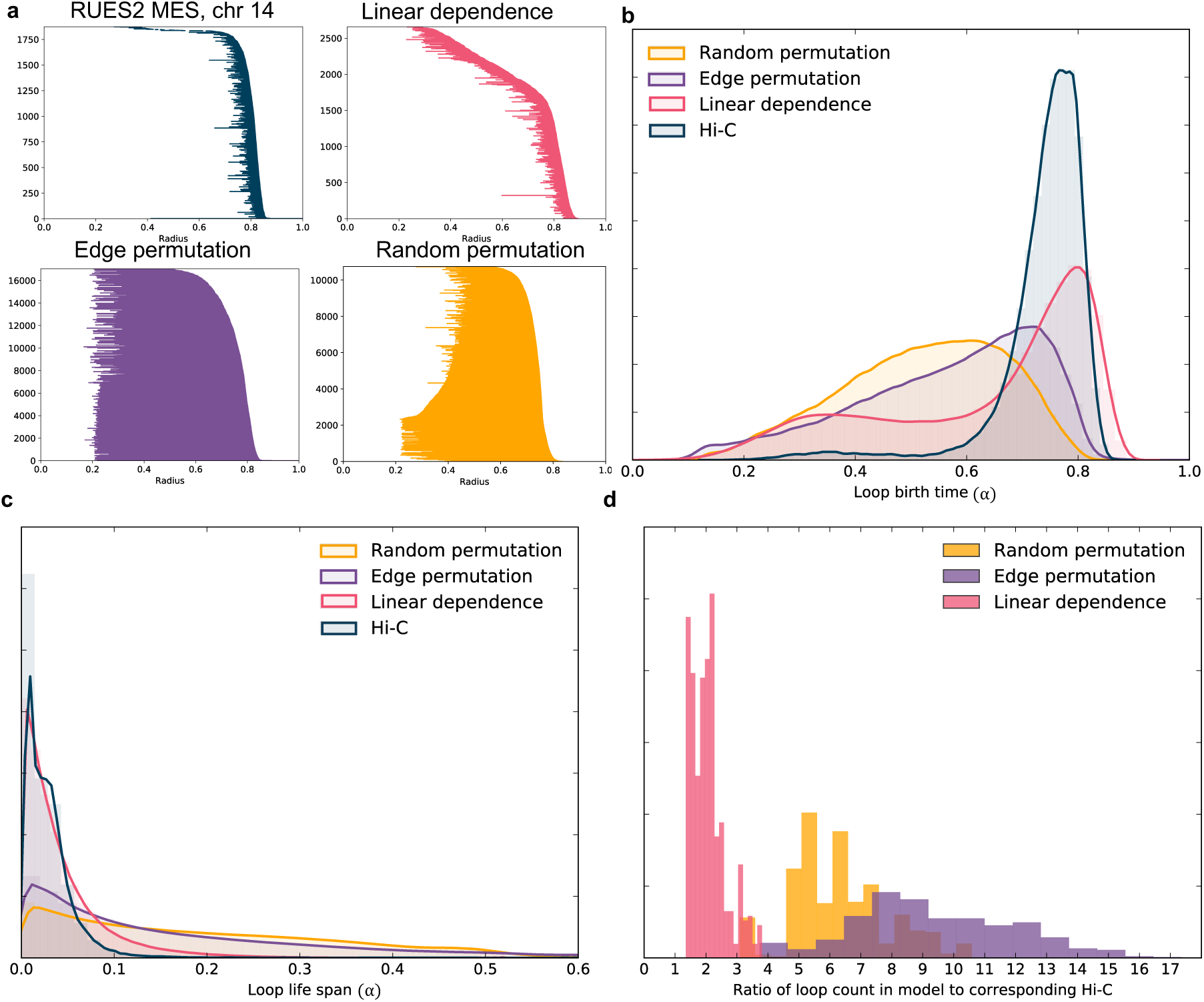
Loop analysis through multiple null models. **(a)** Barcode plots (showing only loop structures) from chromosome 14 of RUES2 MES cells, along with the corresponding barcode plots from the three null models. **(b)** Normalized histogram of the birth times of all loops in RUES2 data from true Hi-C and each null model. Hi-C clearly shows the tightest distribution, with almost no loop birth times before 0.6. **(c)** Normalized histogram of loop life spans (length of a barcode line) shows that the loops from real Hi-C data are very small, similar to the linear dependence model. **(d)** Normalized histogram of the ratios of numbers of loops in each null model to the number of loops in the corresponding Hi-C matrix, showing that all null models result in significantly more loop structures overall than those found in real Hi-C data.

One of the most striking patterns in comparing these null models to the true data is that the loops in Hi-C tend to be born at a much higher value of*α*, and survive only a short time, as noted previously. The model that most closely resembles this pattern is the linear dependence model, but the distribution of birth times shows that the linear dependence model has a long tail towards the earlier birth times which is absent in real Hi-C data (Figure 3b). The short life span, which will generally correspond to smaller loops, is also much more consistent with the linear dependence model, although somewhat more pronounced in Hi-C (Figure 3c). The fact that the linear dependence null model shows these same patterns as Hi-C data suggests that the linear property (nodes that are close together in index, or linear distance, are also close in 3D space) is sufficient to explain the short loops and birth times concentrated at high values of *α*. Long loops appear to be created when nodes with a large difference in their indices (large linear distance) are close together, as demonstrated by the loops with very long life spans in the two permutation models. This appears to be very rare in Hi-C data; the majority of loop interactions we observe are relatively short-lived and involve few genomic bins, consistent with findings from more traditional chromosome loop structures which have been shown to be almost exclusively between loci less than 2Mb apart [25].

The greatest difference between the linear dependence model and Hi-C data can be seen in the number of loops identified (Figure 3d). Although the linear dependence model is again the most similar to real data in this regard, it typically still contains over twice the number of loops as the corresponding Hi-C matrix. This observation could be explained by the existence of topologically associating domains (TADs) or compartments in real chromosomes, which are absent from the linear dependence model but a defining characteristic of Hi-C. These structures serve to isolate chromosome segments from each other, which likely prevents the formation of loops between them. If loops can largely only be formed within a TAD or compartment, this would significantly restrict the total number of feasible loops which could explain the pattern we observe in the persistence diagrams of Hi-C data.

### 3.5 Comparing topologies across differentiation

Although there are some evident changes in topological structures measured by TDA through differentiation, the larger differences exist between the three main cell lines (RUES2, WTC11, and H1, see Figure 4). Interestingly, there does not seem to be any more similarity, measured by bottleneck distance averaged over all chromosomes, between cells at the same stages of differentiation across the three cell lines than cells at different stages of differentiation. For example, the distance value between the two car-diac progenitor samples from RUES2 and WTC11 appears no lower than the value between RUES2 CP and WTC11 CM cells, and H1 ESC and RUES2 ESC seem no more similar to each other than H1 ESC and RUES2 MES or RUES2 ESC and H1 ME cells. Global topological features identified by TDA at the genome-wide scale therefore seem to be determined more by cell line than differentiation stage.

**Figure 4:**
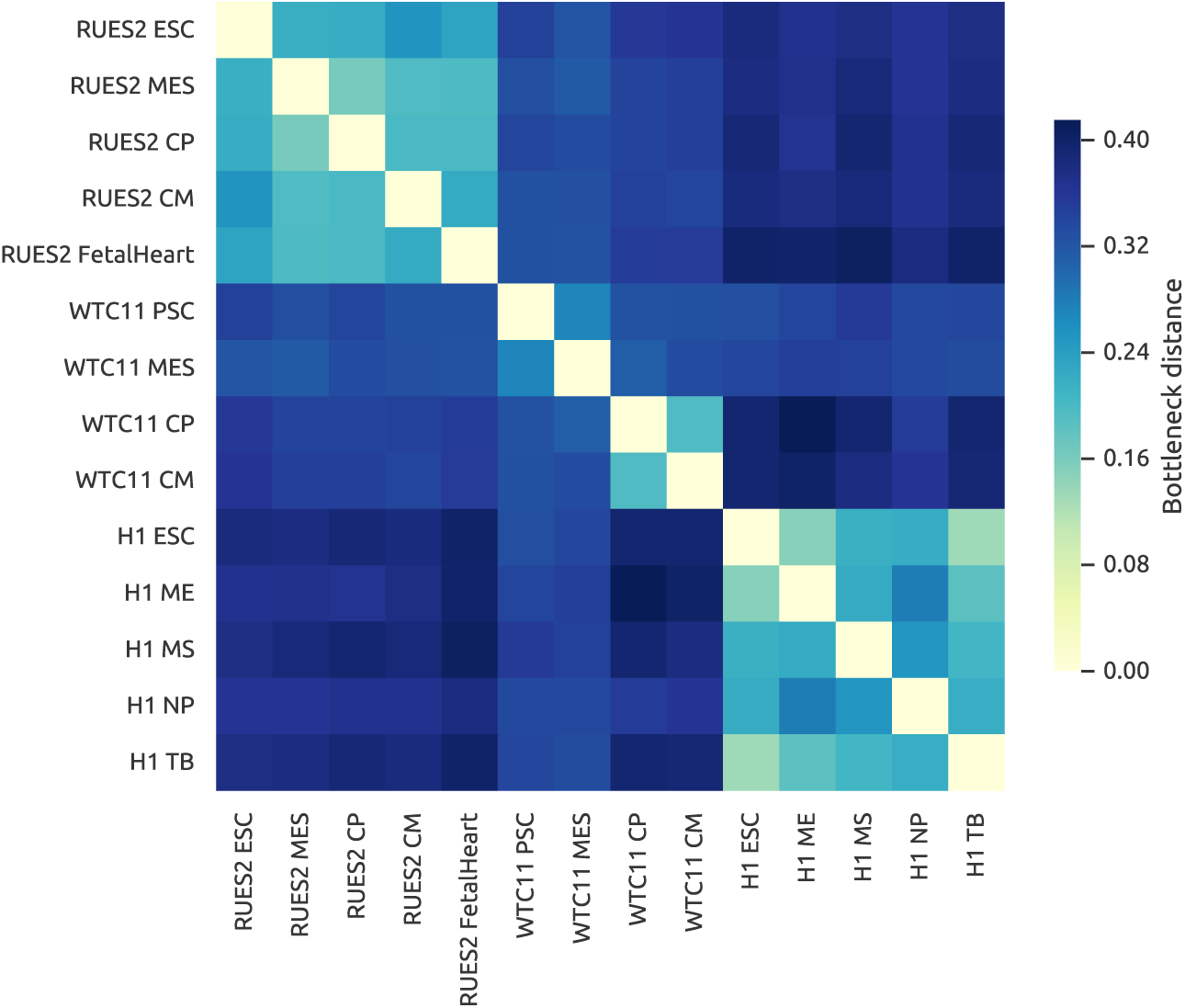
Comparison of all 14 samples studied. This heatmap represents the bottleneck distances between each pair of samples in our data, averaged over all 22 chromosomes. With the exceptions of the high distances between the two later stages and two earlier stages of WTC11 differentiation, the pattern of low distance within one cell line dominates these comparisons. This pattern can be seen by the lighter blocks along the diagonal with limits corresponding to the changes in cell line.

## 4 Discussion

One of the major challenges in the application of TDA to Hi-C data is the computational complexity of the methods combined with the scale of Hi-C. Due to computational limitations, the structures studied here are fairly large-scale; the Hi-C data analyzed is at a relatively low 100kb resolution, and our study only includes 14 samples. By studying smaller sections of the genome, perhaps near a gene of interest or within a particular structure of interest, higher resolution Hi-C or 5C could be used with TDA to identify small-scale topological structures. We were also only able to study intra-chromosomal matrices, but the topology of inter-chromosomal interactions would likely yield interesting insights as well though it would further increase the computational complexity of the problem. Another consequence of the computational complexity of TDA is our focus on only 0- and 1-dimensional features. Emmett et al. (2015) [15] speculate that the 2-dimensional voids may represent interesting biological features such as transcription factories. Future improvements in the data structures and algorithms underlying TDA would significantly improve our ability to study more aspects of the topology of the full genome.

The nature of Hi-C, as an experiment based on cross-linking over a full population, does not permit a transformation from the Hi-C counts to a true distance metric. Our distance matrices therefore do not satisfy the triangle inequality, which may affect the TDA results in unpredictable ways. One possibility to improve this concern is to use any of the methods that estimate a 3D structure from Hi-C, or select only the Hi-C values that satisfy a metric definition [13], and infer distances which would be geometrically consistent. However, this induces another source of error, and it is unclear whether the results would be more reliable.

We have presented the first application of TDA to study the topology of all 22 human autosomal chromosomes. By studying clusters and loops of 14 samples from various cell lines and stages of differentiation, we identify generative principles of chromosome structure. We describe the characteristics of loop structures in chromosomes, the majority of which are very short genomically but with a small number of extremely large outliers suggesting the presence of very long-distance chromosome interactions. Our models suggest that the linearity of the chromosome is sufficient to explain the short lifespan of its loops, but additional structural features specific to Hi-C likely lead to the relatively small number of loops and their late birth times. We also show that topological structure is largely determined by cell line rather than stage of differentiation. TDA shows promise for further analysis of Hi-C data, especially as computational limitations are overcome permitting analysis of higher dimensional features at higher resolution.

## Acknowledgements

The authors would like to thank Alessandro Bertero and William S. Noble for useful information about their data, and Guillaume Marçais for comments on the manuscript.

## S1 Proof of correctness of the loop trace-back algorithm

We will first briefly introduce some concepts related to persistent homology, and then provide a proof that under certain assumptions, our loop trace-back algorithm will find the optimal loop of each homology class.

We restrict all the discussion about homology and persistent homology in this paper over **Z**_2_ field. Let **G** be a simplicial complex, and **G**_*k*_ be the set of simplices of **G** with dimension *k*. A *k*-chain *c* on **G** is defined as the sum of *k*-simplices, 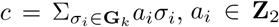. The boundary of a *k*-simplex is the sum of its (*k−*1)-faces, and the boundary of a *k*-chaincan be defined as the sum of all boundaries of its *k*-simplex components. We can then define a boundary map as *∂*_*k*_:*C*_*k*_(**G**) *→C*_*k−*1_(**G**), where *C*_*k*_(**G**) is a vector space formed by all the *k*-chains on **G**. We define the kernel of *∂*_*k*_ as *Z*_*k*_(*G*) and the image of *∂*_*k*+1_ as *B*_*k*_(*G*). It can be proved that *B*_*k*_(*G*) is the subgroup of *Z*_*k*_(*G*) [9], and since it is easy to see that *Z*_*k*_(*G*) is an abelian group, *B*_*k*_(*G*) is also a normal subgroup. Hence we can define the homology group, *H*_*k*_(*G*) as the quotient group between *Z*_*k*_(*G*) and *B*_*k*_(*G*), *H*_*k*_(*G*):=*Z*_*k*_(*G*)/*B*_*k*_(*G*).

As described in Methods, the persistent homology algorithm involves building a series of VR complexes by varying the parameter *α*, creating different homology classes that appear and disappear at different values of *α*. Let *h*_1*i*_(*α*_0_), *i ∈ {*1, 2, 3, .., *n}* be the i-th one-dimensional homology class formed at *α*_0_, *n* isthetotalnumberofnewlyformed one-dimensional homology classes at the point *α*_0_. We would like to identify a representative element *z ∈ h*_1*i*_(*α*_0_) for each *i*.

For a given parameter *α*_0_, we construct a weighted graph (*K*(*α*_0_), *V, E*) where vertices are all points in *X*, the given data set, and an edge, with weight equal to the distance between the vertices, exists if the weight is not larger than *α*_0_.We denote the weight of an edge *e ∈ E* as *w*(*e*) and denote the weight of any cycle *l* as the sum of all the edges it contains, *w*(*l*)= Σ_e ∈ *l*_ *w*(*e*).

For the graph (*K*(*α*_0_), *V, E*), it is easy to see that *Z*_1_ (*K*(*α*_0_)) is composed of all the cycles on *K*(*α* _0_) *B*_1_ (K (*α* _0_)) is the subset of cycles and *H*_1_ (*K* (*α* _0_)) is the homology group of which each element is the set of cycles with the same homology class. For any cycle *l* ∈ *Z*_1_ (*K* (*α* _0_)), *l* = {*e*_1_, …, *e*_*j*_}, *e*_*i*_ ∈ *E*, we define 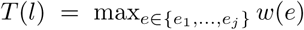 as the point when the cycle is formed. We also denote [*l*] as the corresponding homology class of *l* in *H*_1_ (*K*(*α* _0_)), [*l*] ={*l* + *b|b ∈ B*_1_(*K*(*α*_0_))}, and for any homology class *h ∈ H*_1_(*K*(*α*_0_)), define *T* (*h*) = min_*k ∈ h*_ *T* (*k*) as the point when the homology class appears.

Assuming only one homology class is formed at any given *α*, we want to find its optimal cycle L, defined as:

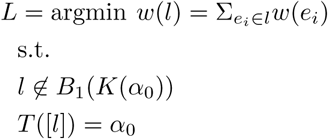

Let *m* be the number of edges in *K*(*α*_0_) with weight *α*_0_, we have the following theorems.

### Theorem 1.

*For a given parameter α*_0_, *if m* = 1, *then any cycle l on K*(*α*_0_) *satisfying the two conditions l* ∉ *B*_1_(*K*(*α*_0_)) *and T* ([*l*]) = *α*_0_ *contains the edge* [*a, b*], *where w*([*a, b*]) = *α*_0_.

*Proof*. Assume for contradiction that there exists a cycle *l* = {*e*_1_, *e*_2_, …, *e*_*p*_} on *K*(*α*_0_), *l* ∉ *B*_1_(*K*(*α*_0_)) and *T* ([*l*]) = *α*_0_, which does not include the edge [*a, b*]. We assume there is only one edge with weight *α*_0_, so *w*(*e*_*i*_) *< α*_0_ for all *e*_*i*_ ∈ {*e*_1_, …, *e*_*p*_}. Then 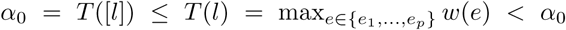, which contradicts the first statement.

### Theorem 2.

*For a given parameter α*_0_, *if m* = 1, *then any cycle l which belongs to a homology class h satisfying the conditions h* ∈ *H*_1_(*α*_0_) *and T* (*h*) = *α*_0_ *contains the edge* [*a, b*] *with w*([*a, b*]) = *α*_0_.

*Proof*. Assume for contradiction that there exists a cycle *l* ∈ *h, l* = {*e*_1_, *e*_2_, …, *e*_*p*_}, that does not include [*a, b*], and *h ∈ H*_1_(*α*_0_) and *T* (*h*) = *α*_0_.There is only one edge with the weight *α*_0_, so *w*(*e*_*i*_) < *α*_0_*∀e*_*i*_ ∈ {*e*_1_, …, *e*_*p*_}.Then 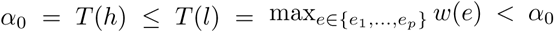, which is a clear contra-diction.

### Theorem 3.

*For a given parameter α*_0_, *if m* = 1 *and there is one homology class appearing at α*_0_, *then any cycle l on K*(*α*_0_) *which contains the edge* [*a, b*], *w*([*a, b*]) = *α*_0_, *does not belongs to B*_1_(*K*(*α*_0_)).

*Proof*. Assume ∃*l* = {*e*_1_, *e*_2_, …, *e*_*p*_} + [*a, b*], *e*_*i*_ ∈ *E*, such that *l* ∈ *B*_1_(*K*(*α*_0_) Since there is one homology class appearing at *α*_0_, ∃h ∈ *H*_1_(*K*(*α*_0_)) such that *T*(*h*) = *α*_0_. Choose one cycle *l*′ ∈ *h*, from Theorem 2, we know that *l*′ contains the edge [a; b], then we can write *l*′ as 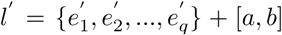. Since 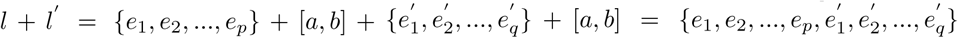 still belongs to the homology class *h*. Then 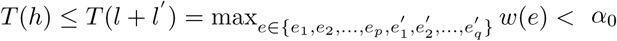, which is again a clear contradiction.

From Theorem 3, we can see that for any cycle *l* on the graph *K*(*α*_0_) which contains the edge [*a, b*] with the weight *w*([*a, b*]) = *α*_0_, [*l*] = {*l* + *b|b* ∈ *B*_1_(*K*(*α*_0_))} is an non identity element in *H*_1_(*K*(*α*_0_)) and *T* ([*l*]) = *α*_0_.

For a given parameter *α*_0_, if *m* = 1, let [*a, b*] bethe edge with the weight *α*_0_, remove [*a, b*] from the graph *K*(*α*_0_) and let *p*_*short*_ be the minimum weight path from *a* to *b* on that new graph, according to Theorem 3, we know that if there is one homology class appearing at *α*_0_, [*p*_*short*_ + [*a, b*]] ∉ *B*_1_(*K*(*α*_0_)) and *T* ([*p*_*short*_ + [*a, b*]]) = *α*_0_. According to Theorem 1, we know that any cycle *l* on *K*(*α*_0_) satisfying the conditions *l* ∉ *B*_1_(*K*(*α*_0_)) and *T* ([*l*]) = *α*_0_ contains the edge [*a, b*], therefore we have *w*(*p*_*short*_ + [*a, b*]) ≤ *w*(*l*). Then the cycle *p*_*short*_ + [*a, b*] is the optimal cycle as previously defined.

**Figure S1:**
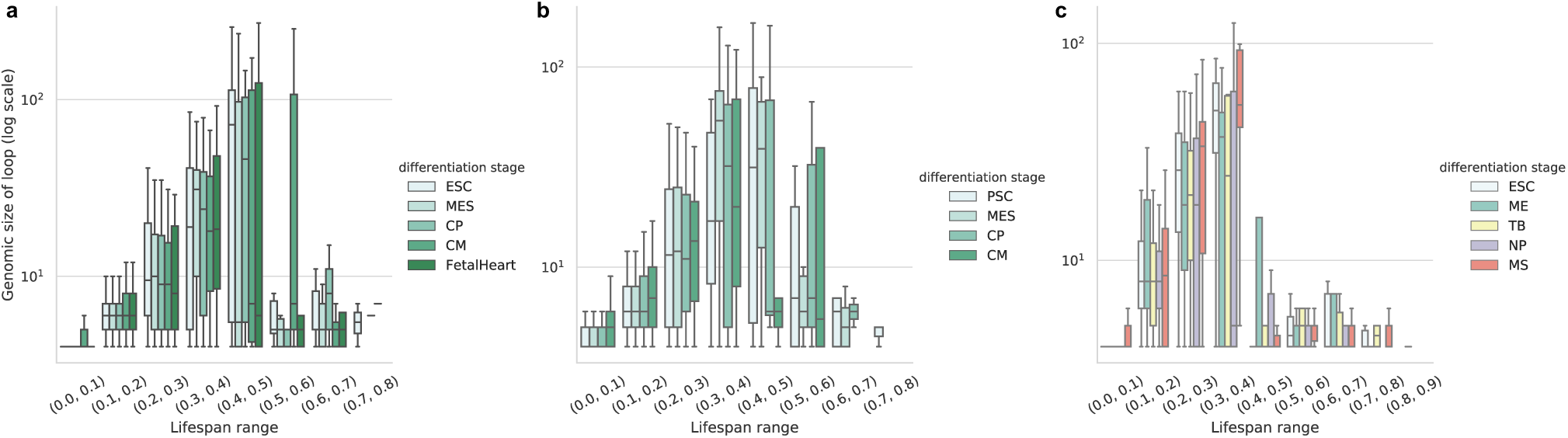
Distributions of genomic loop sizes across loop lifespans, separated by differentiation stage. The log-scaled y-axis again represents the total number of genomic bins returned by the loop traceback method. (a) RUES2 cell line (b) WTC11, and (c) H1. Note that because the H1 data is not a linear trajectory through the differentiation process like the RUES2 and WTC11 data, this is represented by a different color scheme.

**Figure S2:**
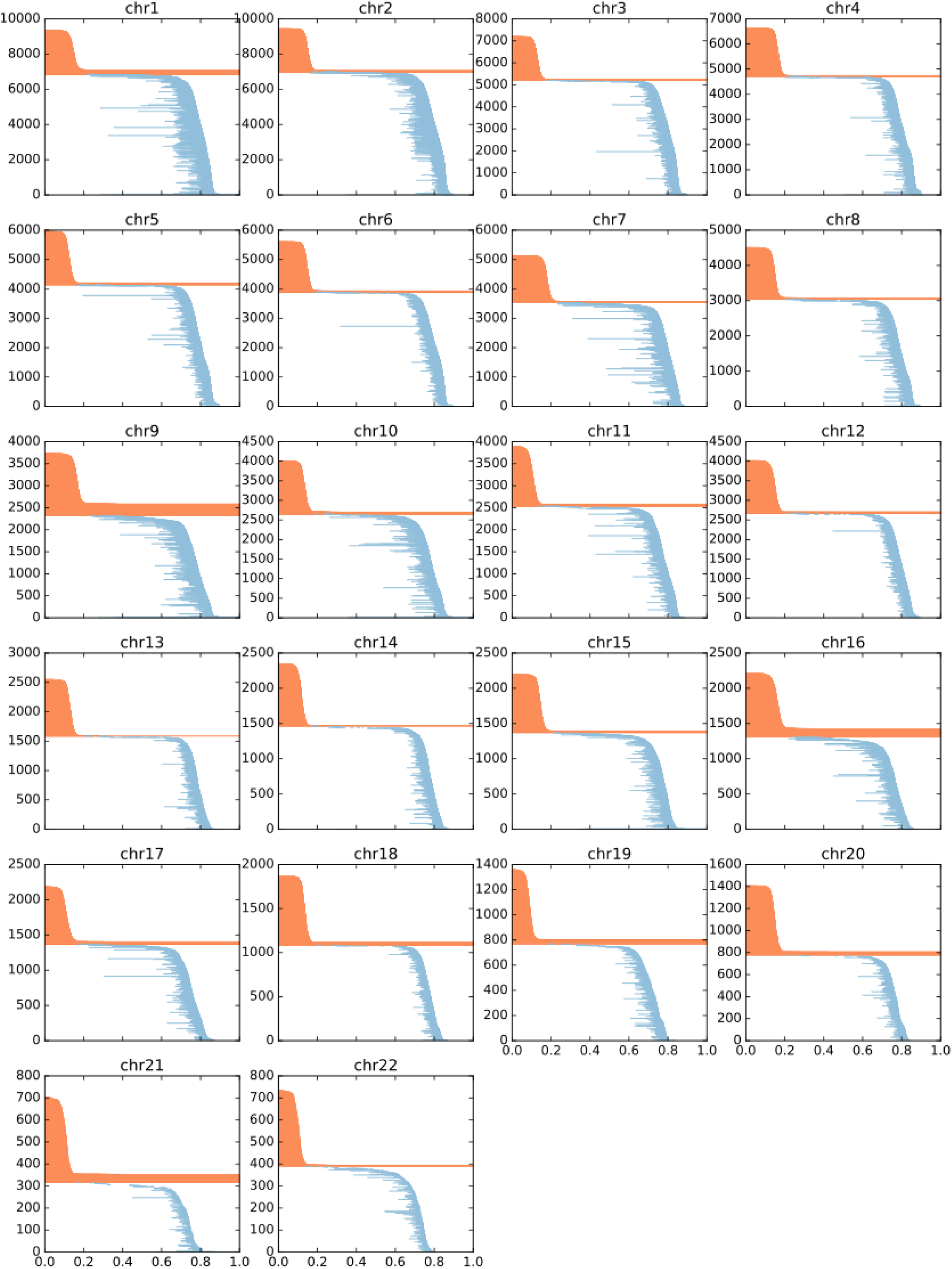
RUES2 ESC barcode diagrams

**Figure S3:**
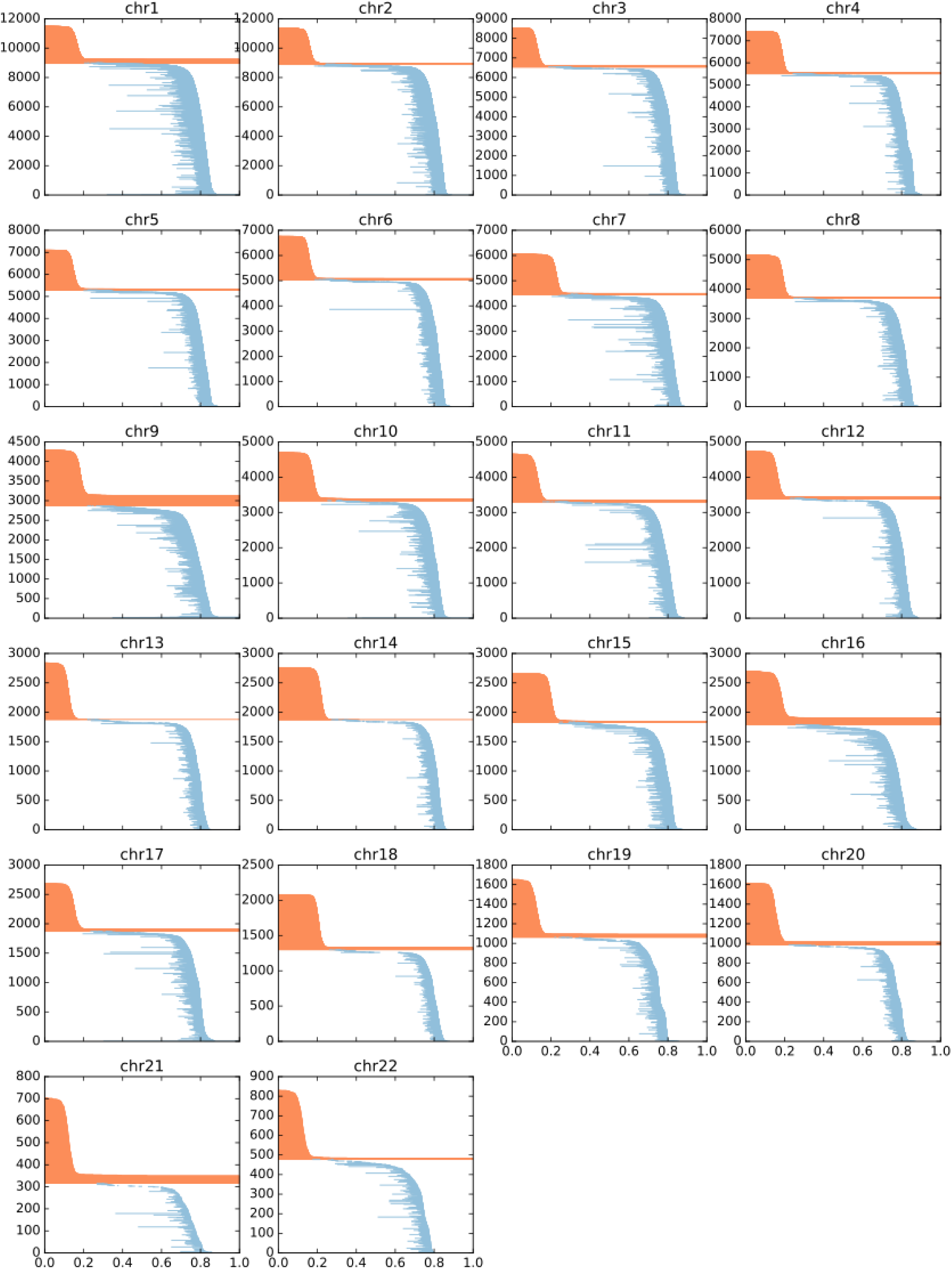
RUES2 MES barcode diagrams

**Figure S4:**
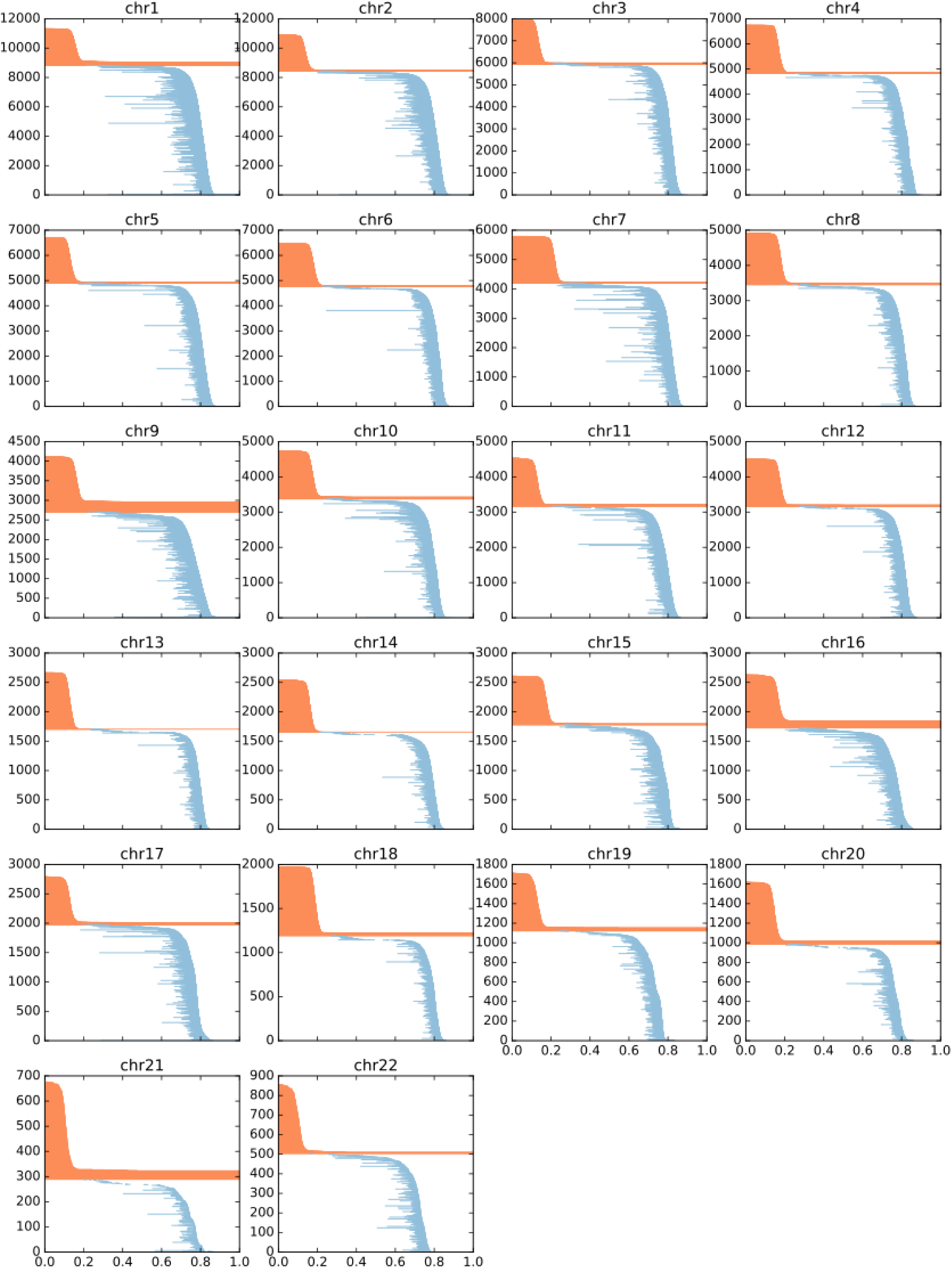
RUES2 CP barcode diagrams

**Figure S5:**
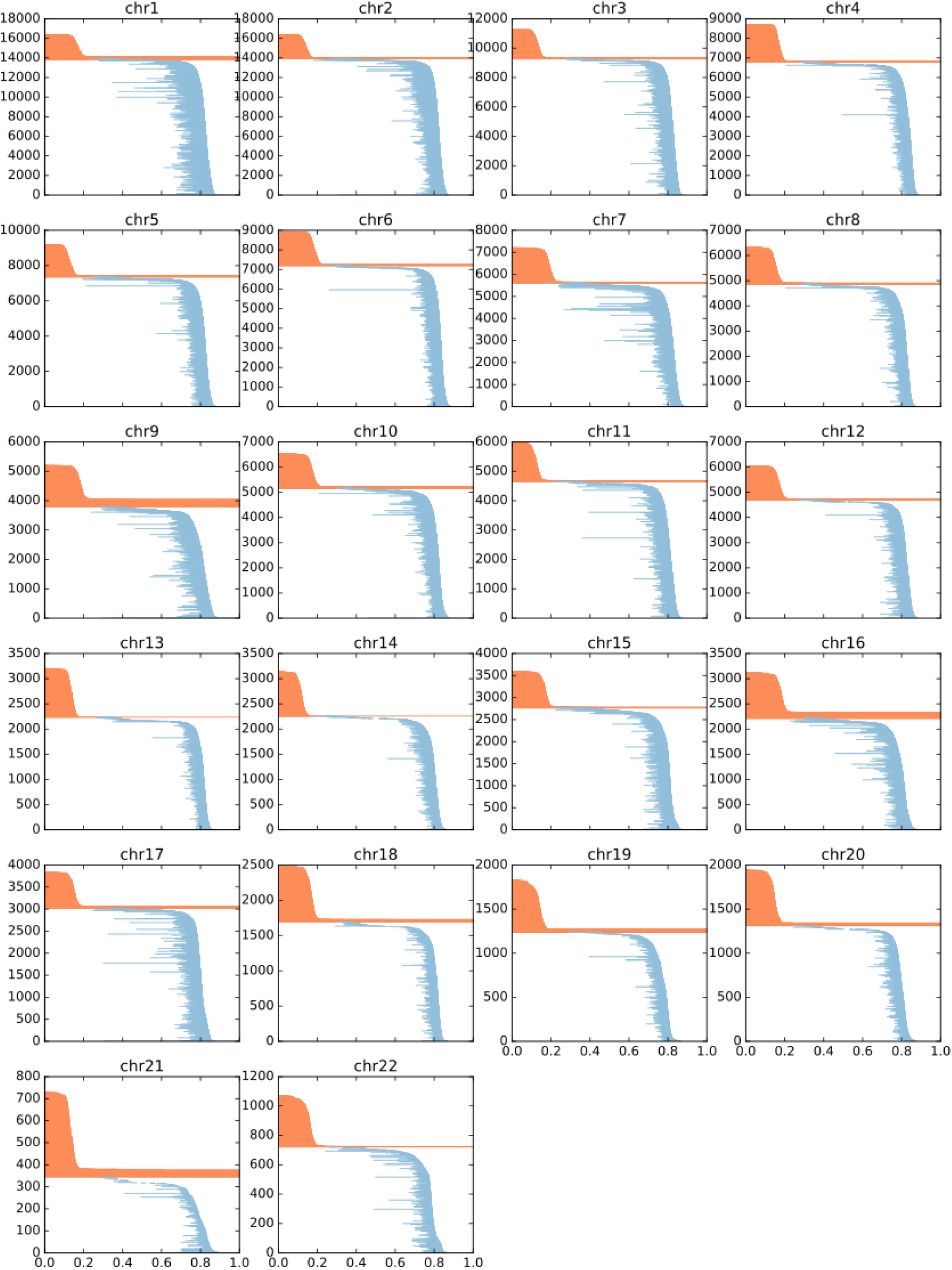
RUES2 CM barcode diagrams

**Figure S6:**
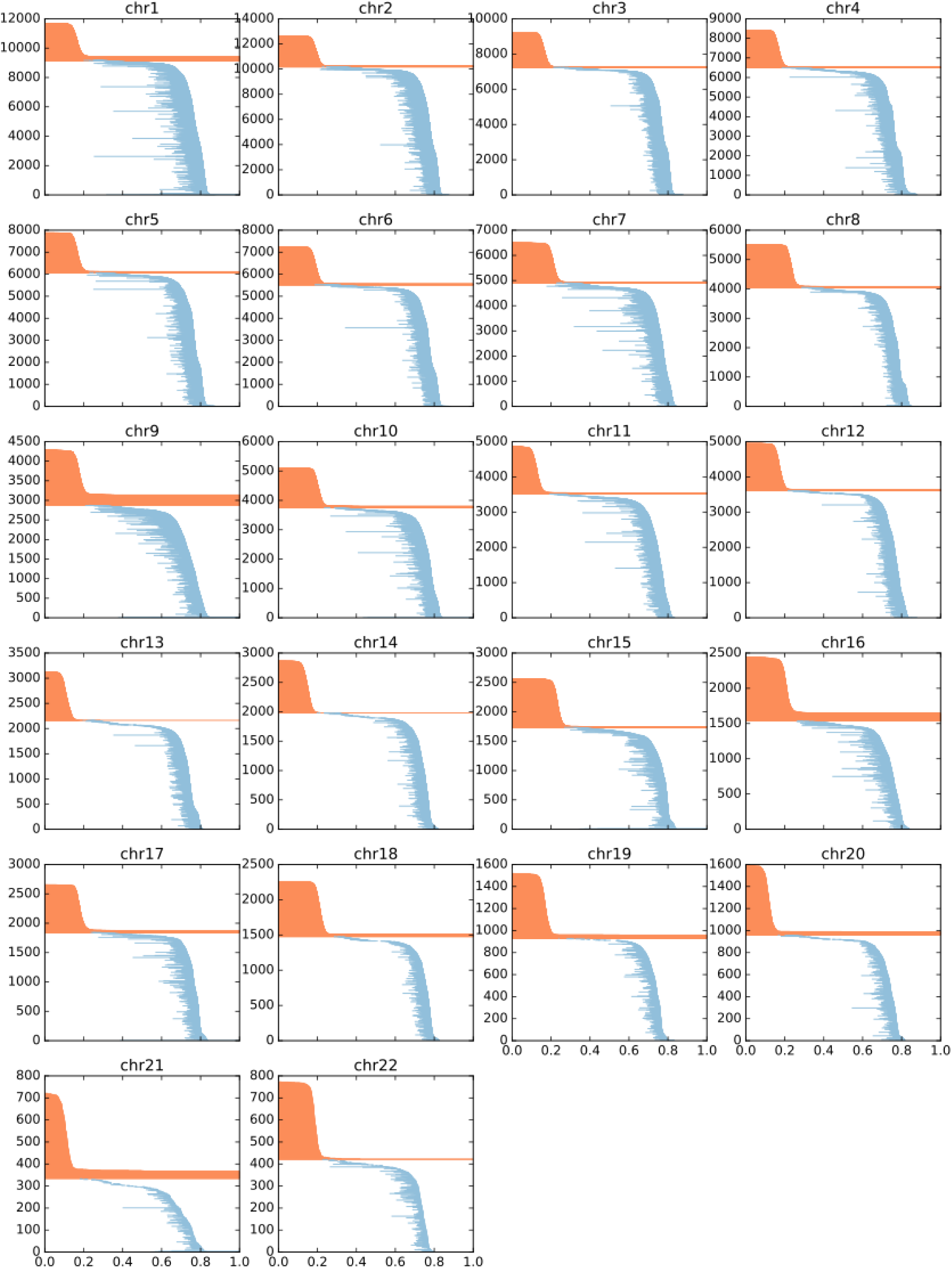
RUES2 Fetal Heart barcode diagrams

**Figure S7:**
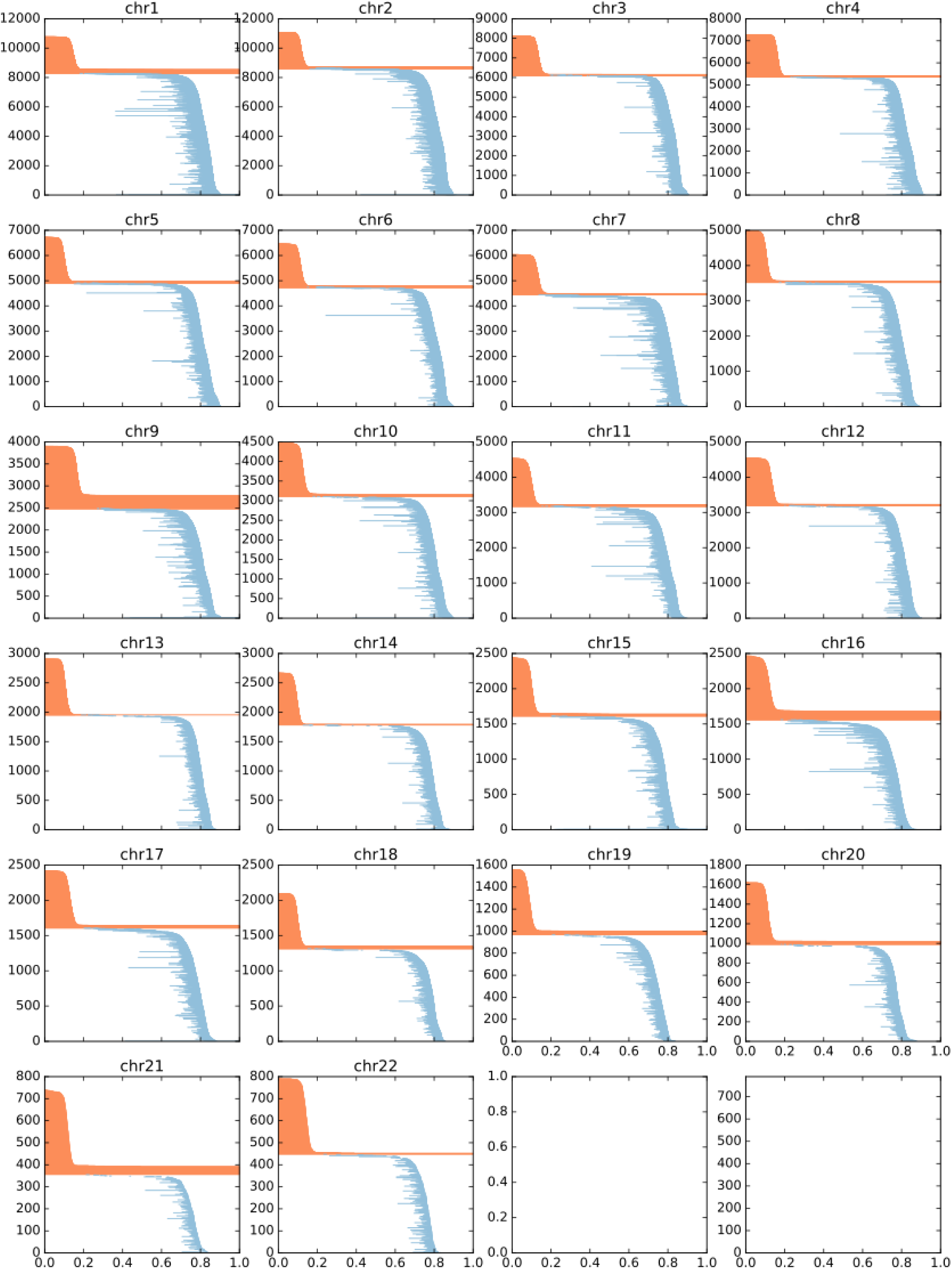
WTC11 PSC barcode diagrams

**Figure S8:**
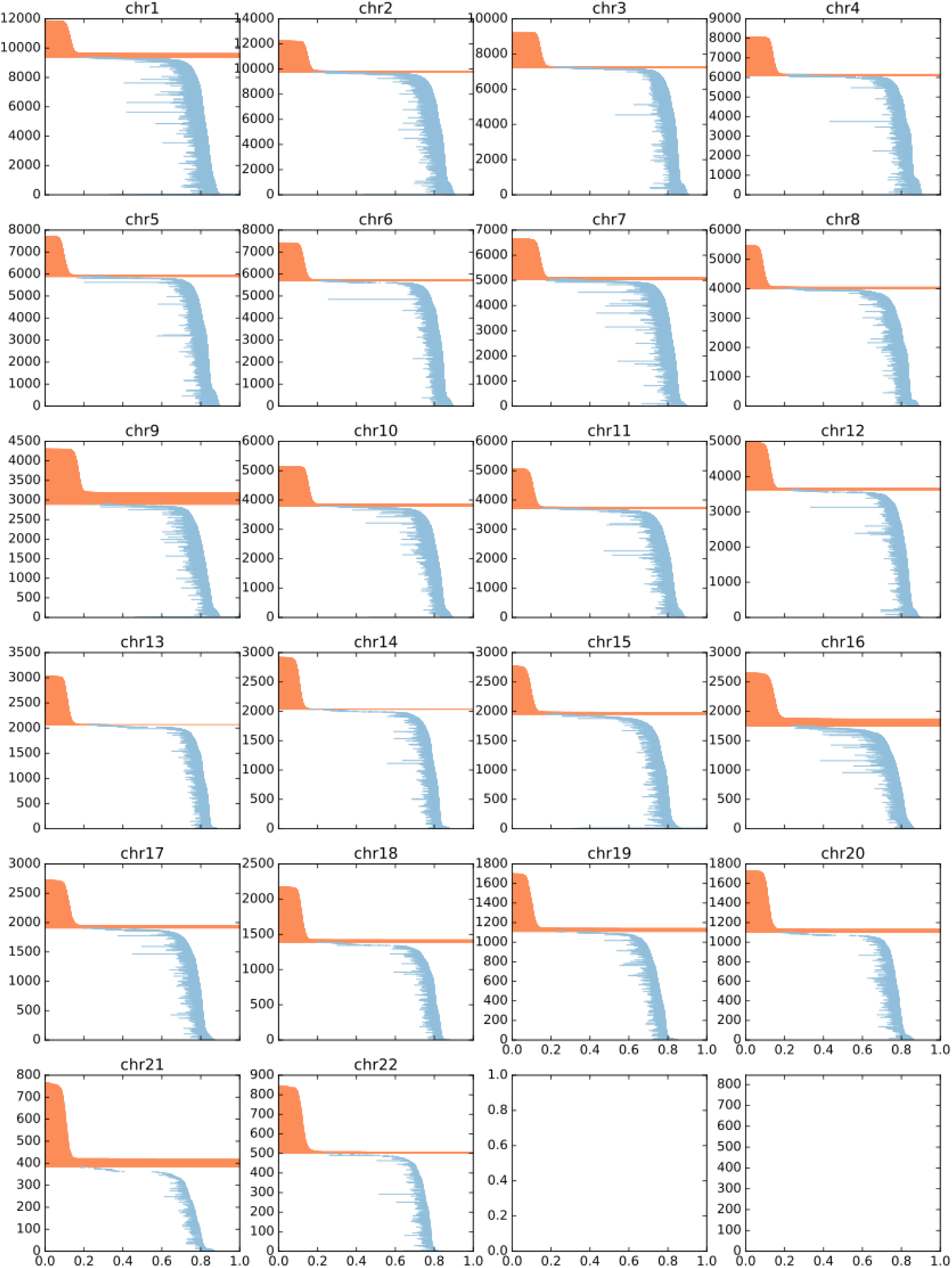
WTC11 MES barcode diagrams

**Figure S9:**
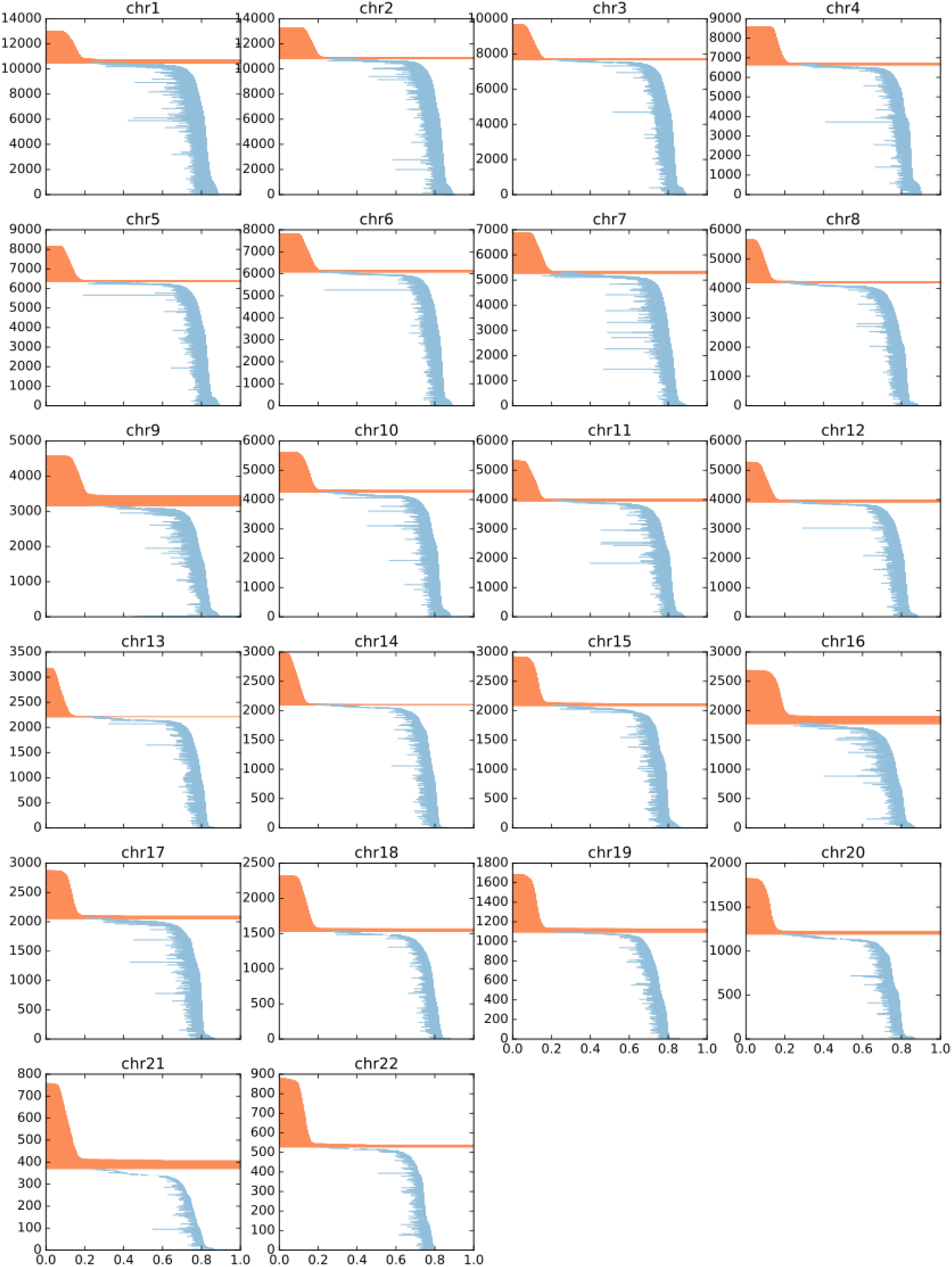
WTC11 CP barcode diagrams

**Figure S10:**
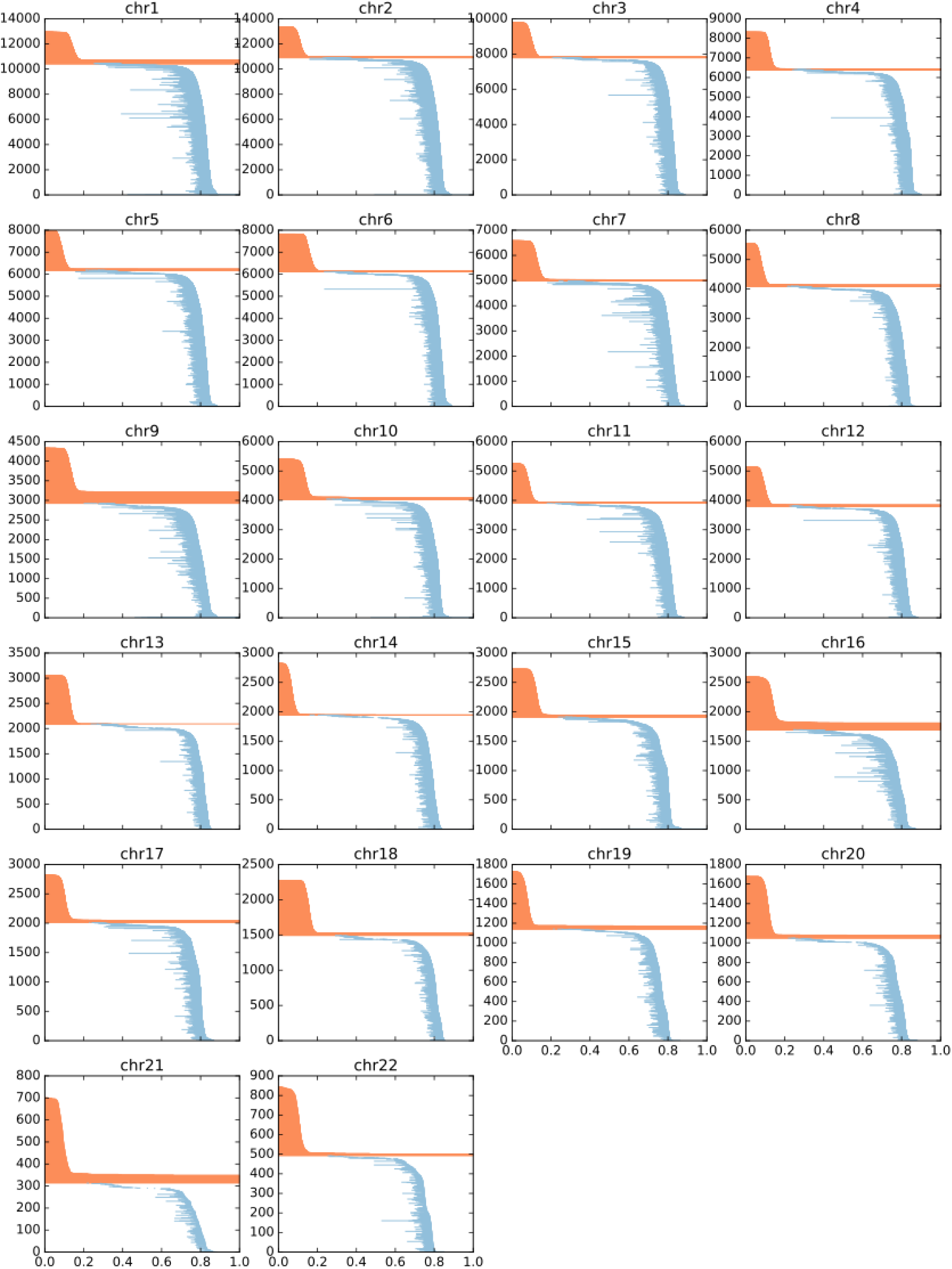
WTC11 CM barcode diagrams

**Figure S11:**
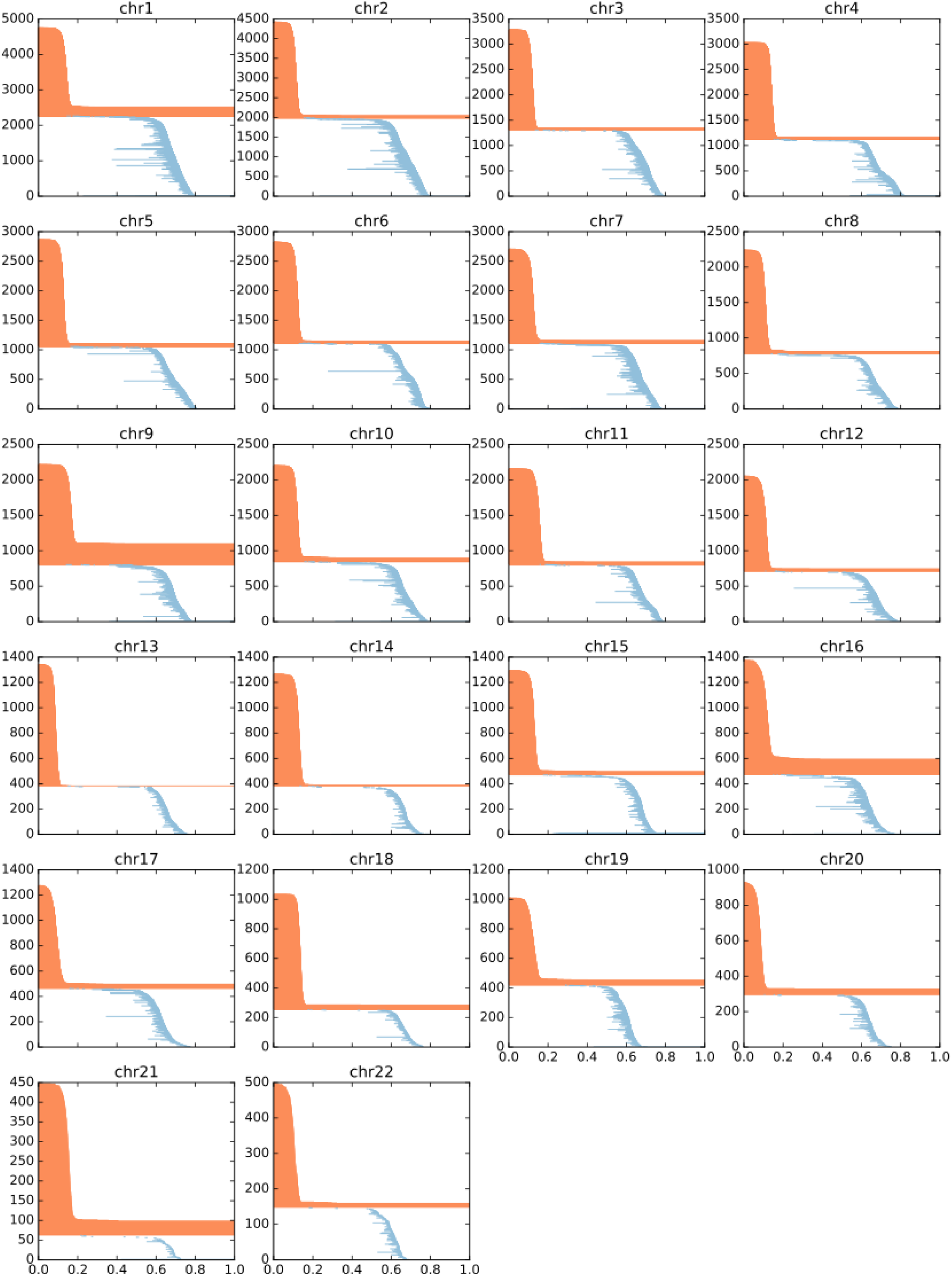
H1 ESC barcode diagrams

**Figure S12:**
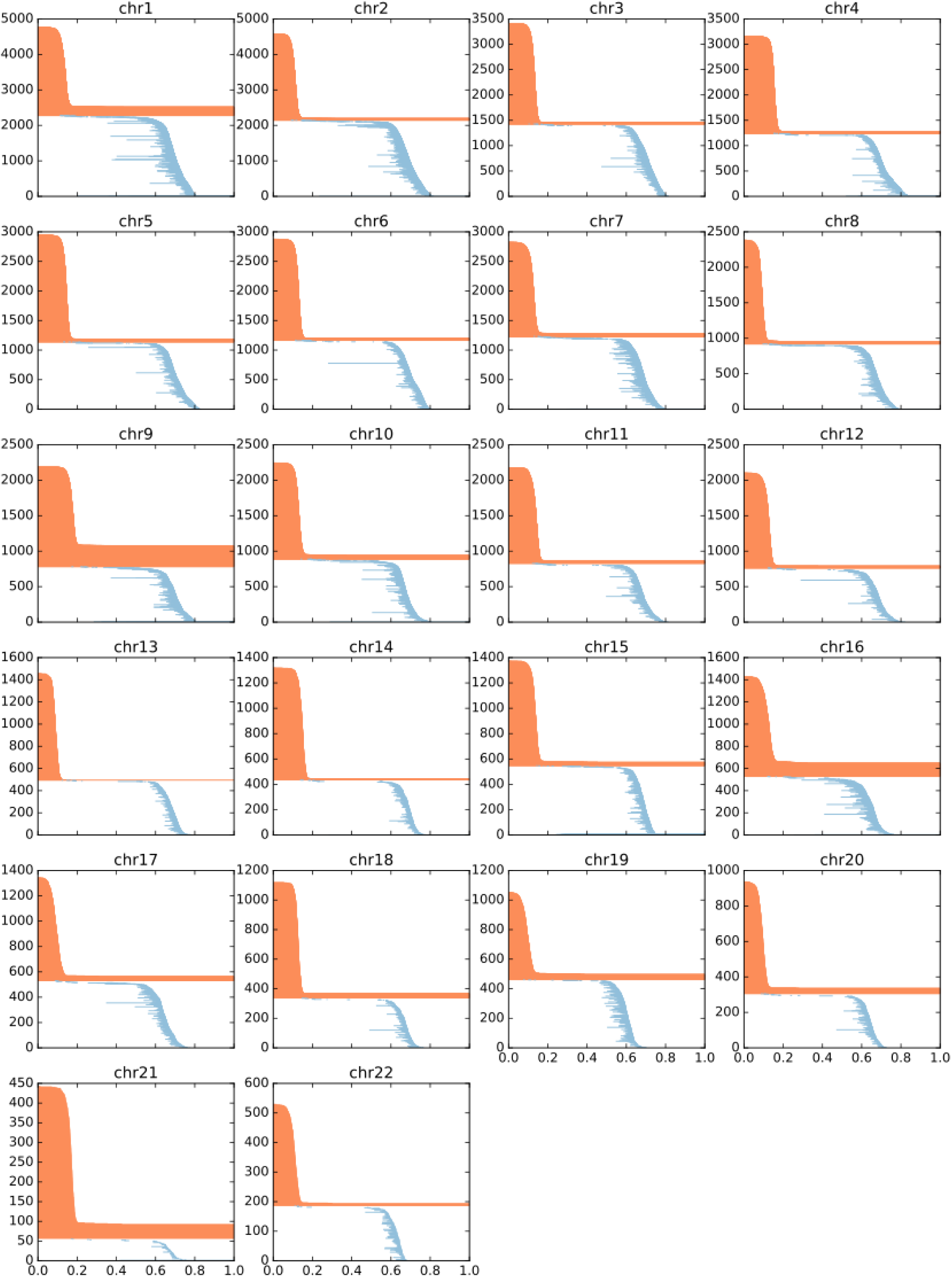
H1 ME barcode diagrams

**Figure S13:**
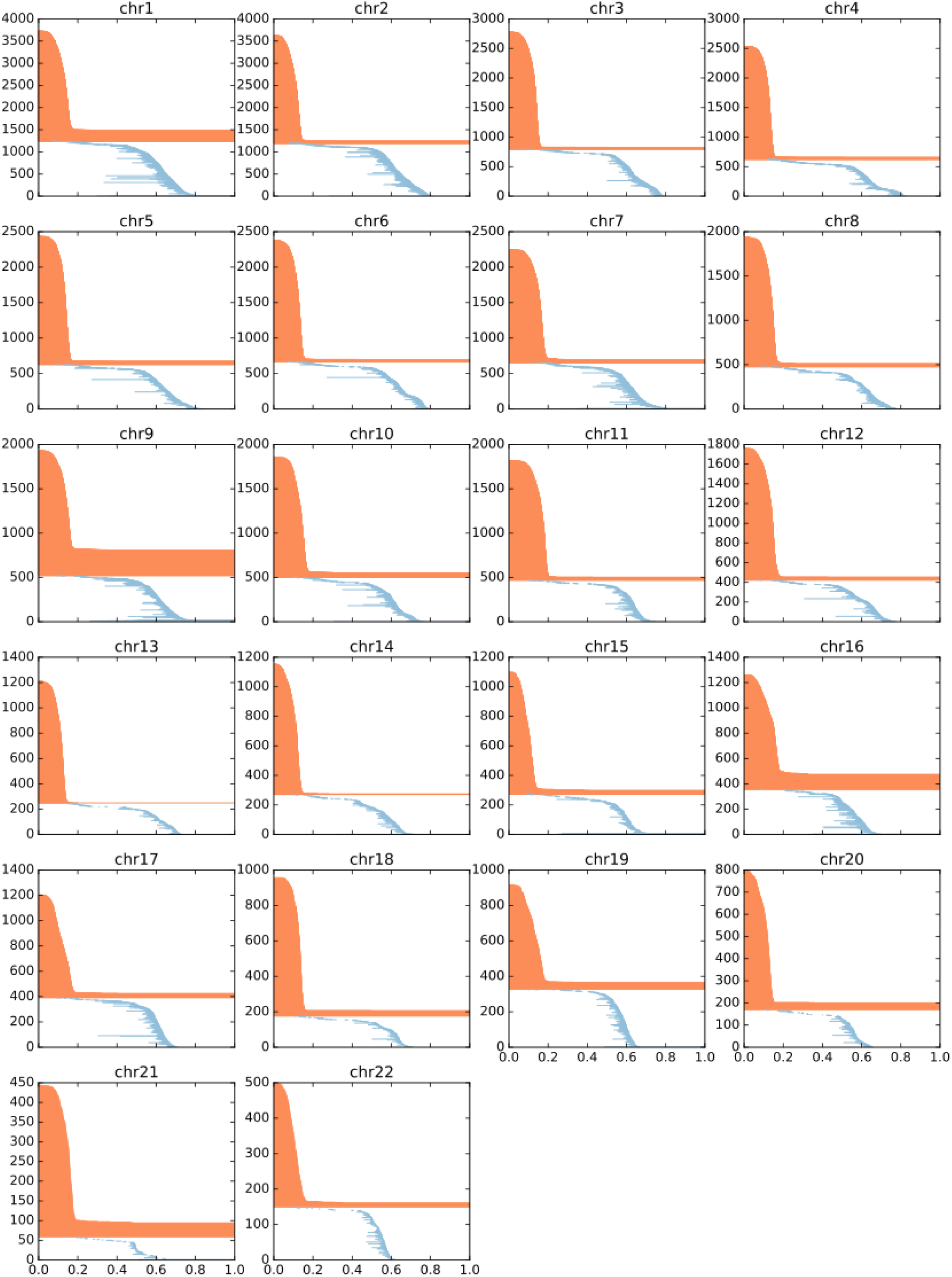
H1 MS barcode diagrams

**Figure S14:**
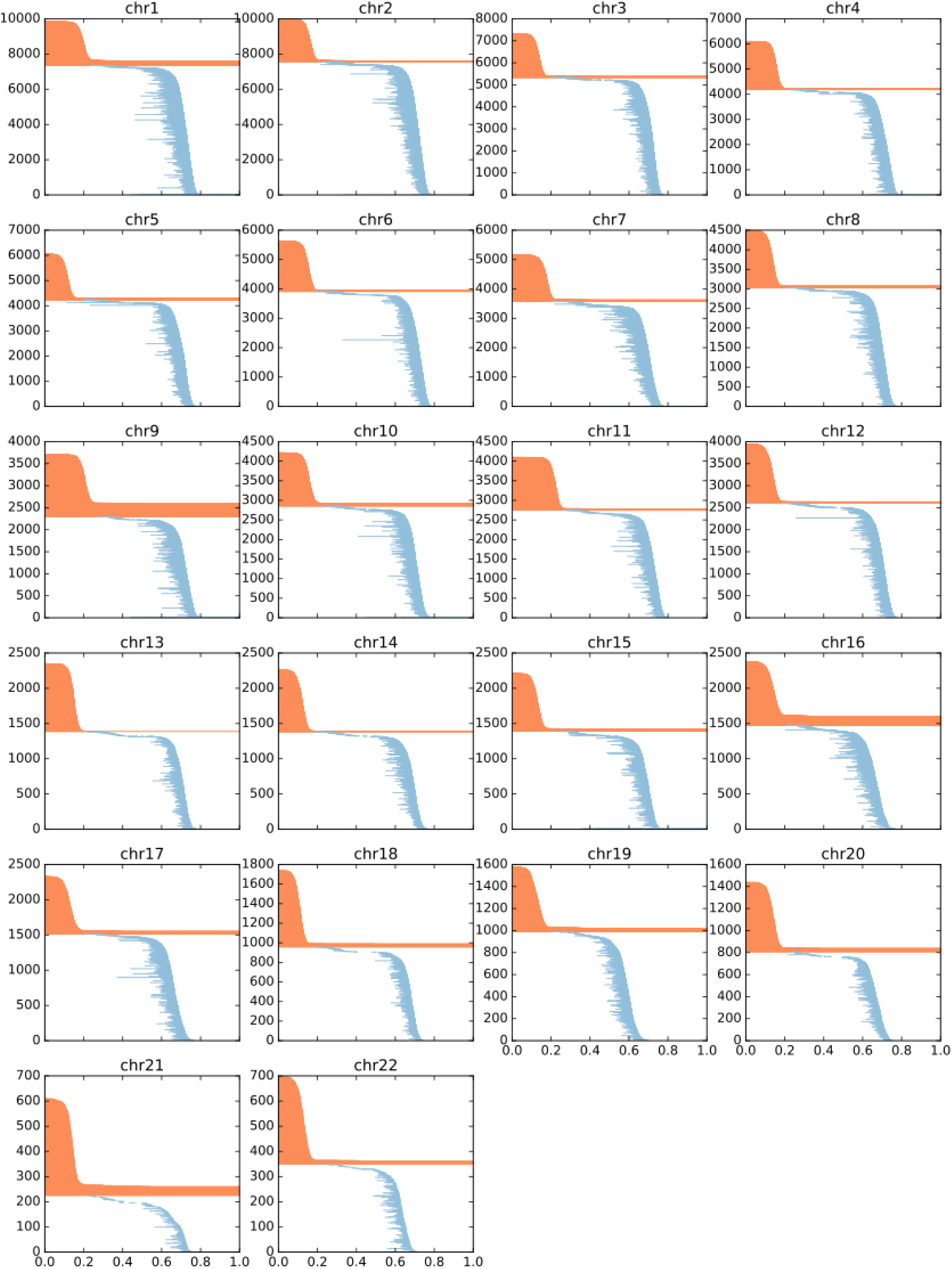
H1 NP barcode diagrams

**Figure S15:**
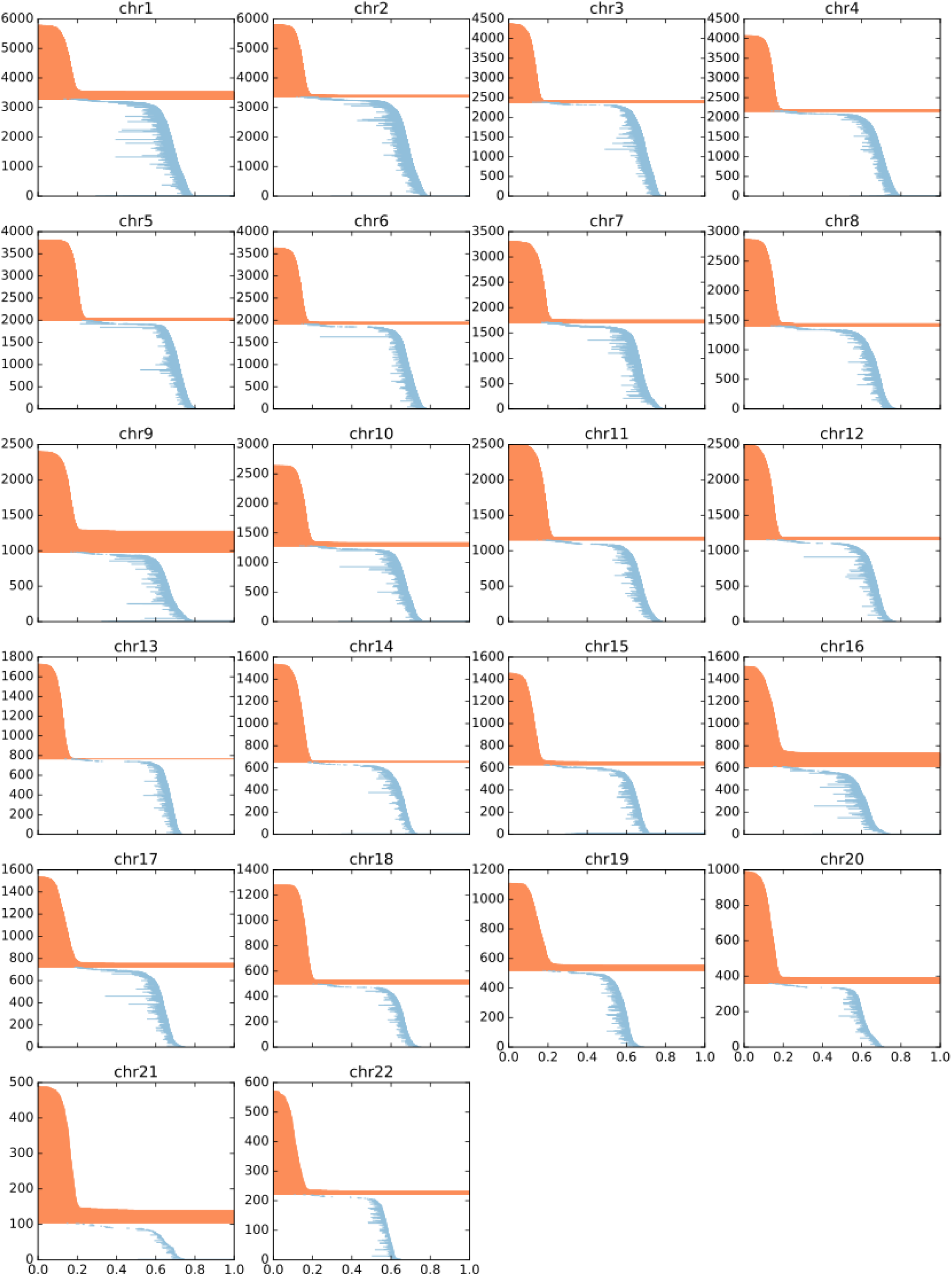
H1 TB barcode diagrams

**Figure S16:**
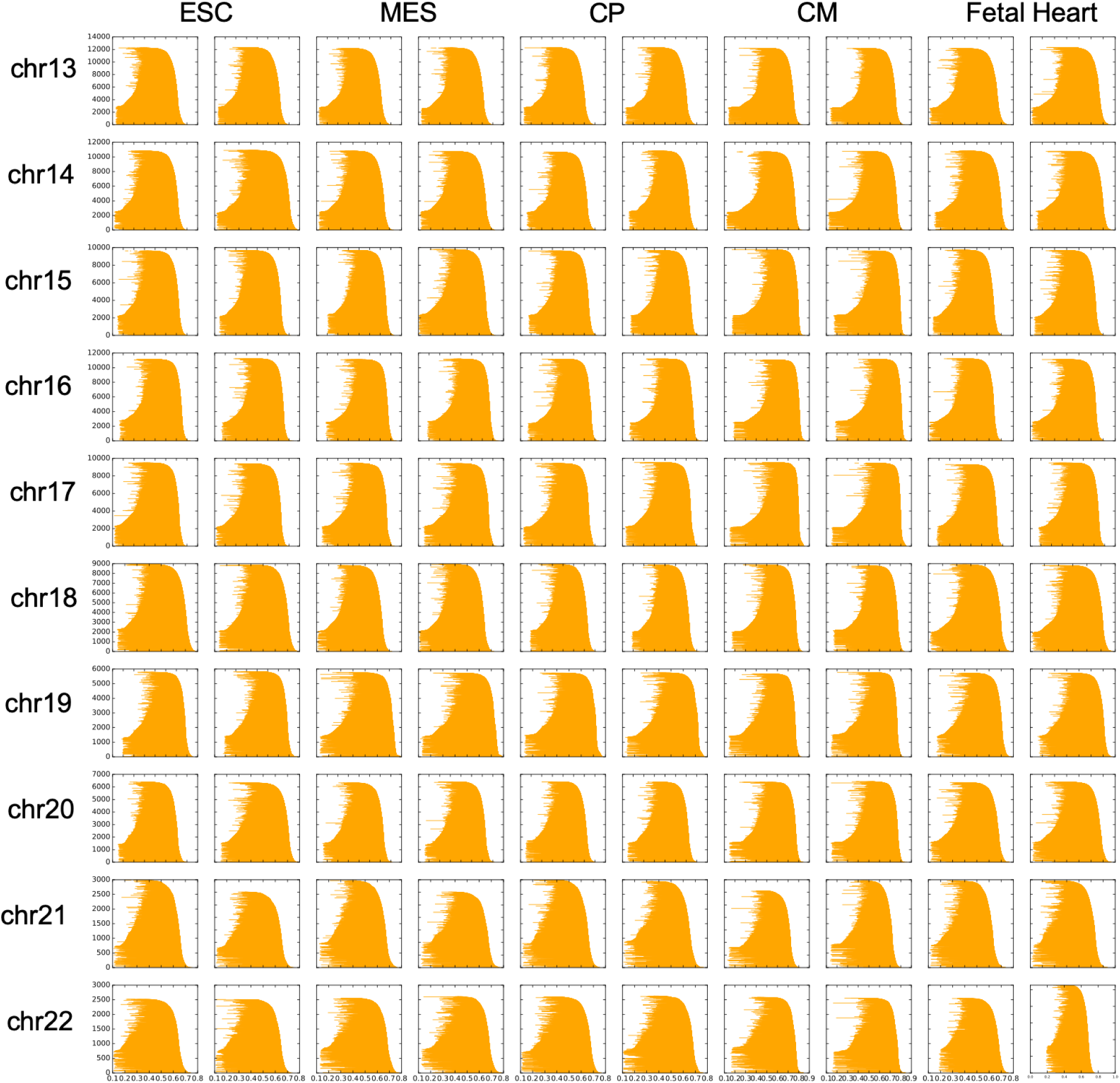
Barcode diagrams for the random permutation null model. Two random examples were chosen from the ten generated for each cell type.

**Figure S17:**
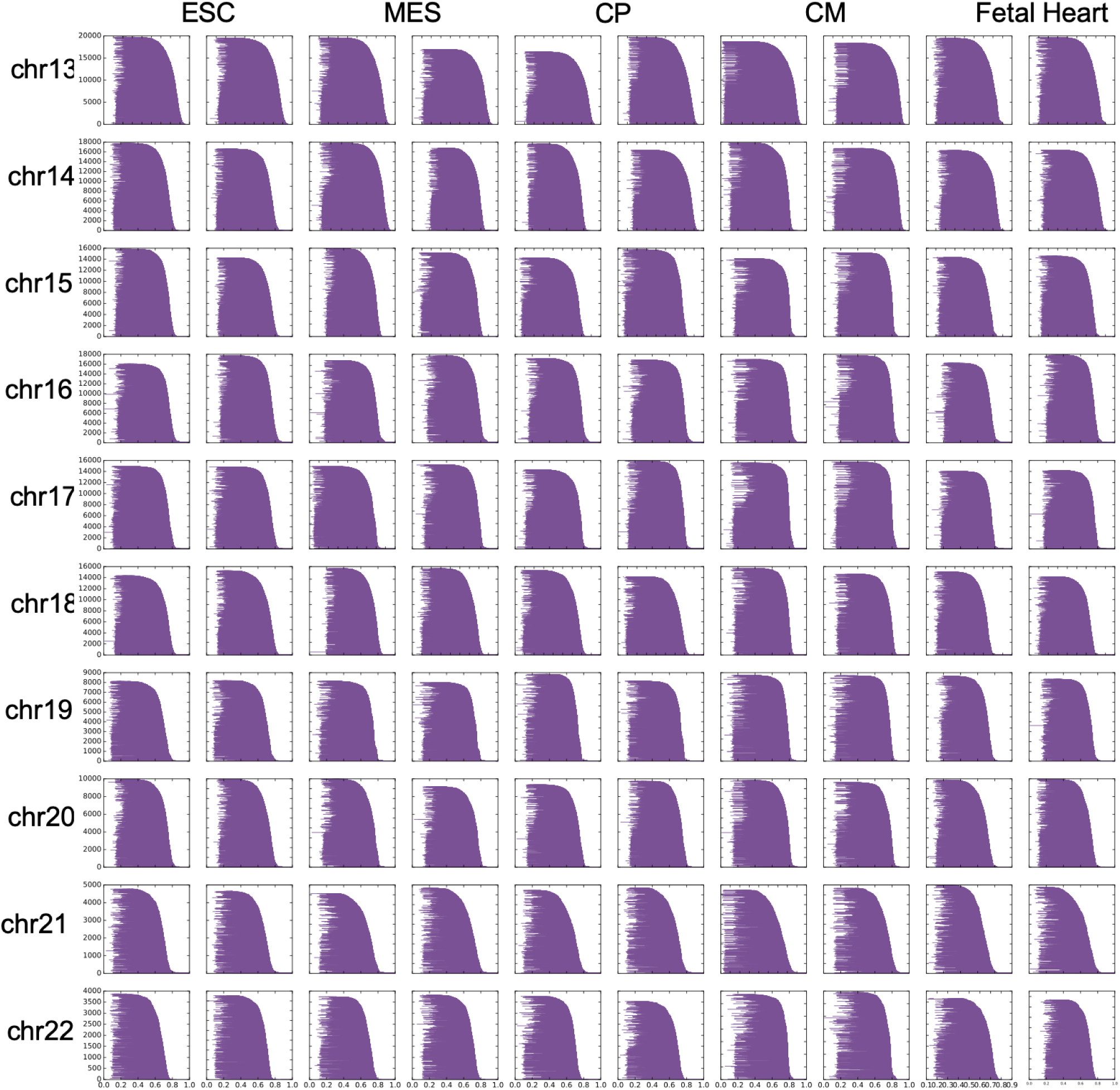
Barcode diagrams for the edge permutation null model. Two random examples were chosen from the ten generated for each cell type.

**Figure S18:**
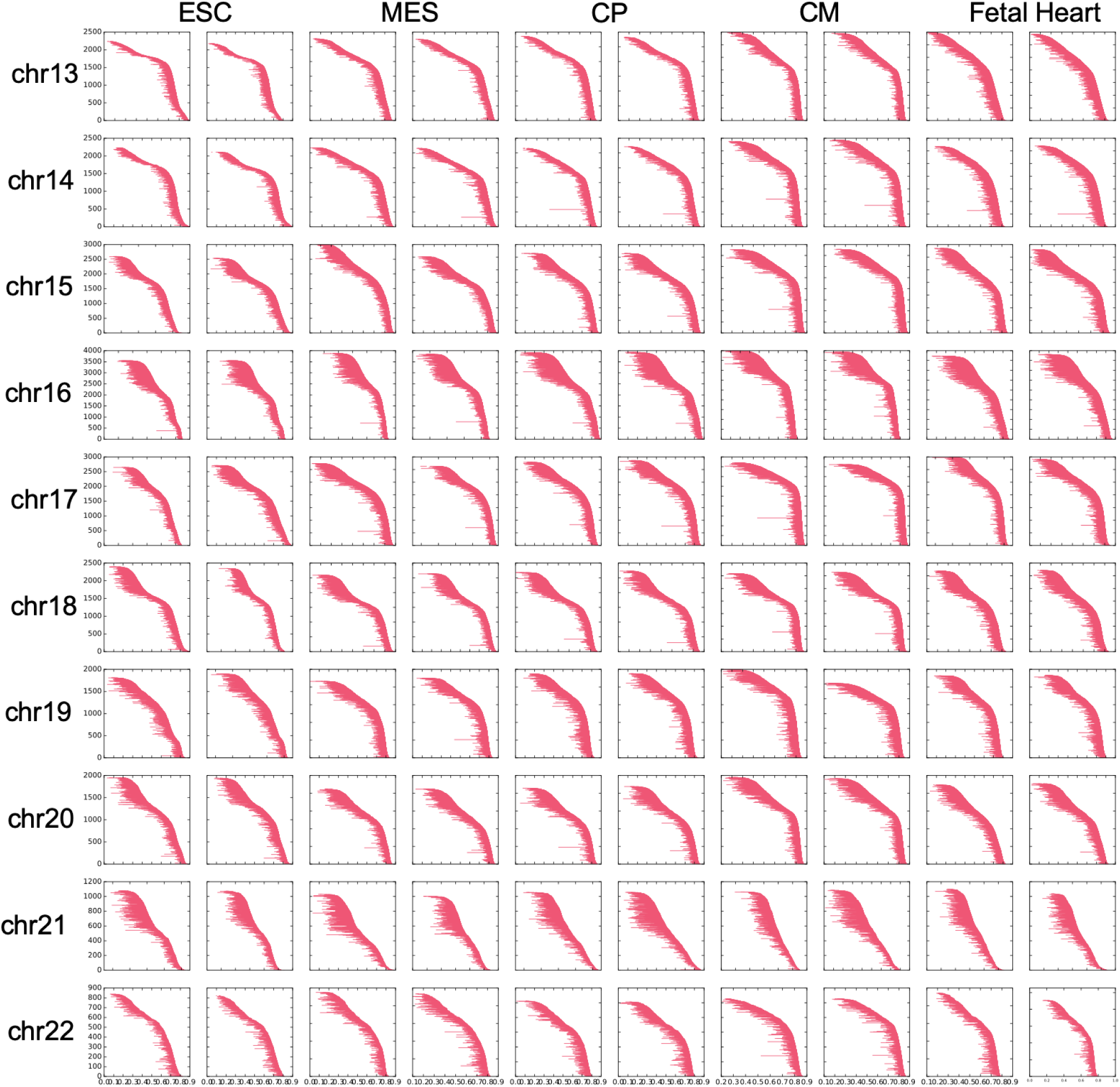
Barcode diagrams for the linear dependence null model. Two random examples were chosen from the ten generated for each cell type.

## References

[1] Javier Arsuaga, Tyler Borrman, Raymond Cavalcante, Georgina Gonzalez, and Catherine Park. Identification of copy number aberrations in breast cancer subtypes using persistence topology. Microarrays, 4(3):339–369, 2015.

[2] Ferhat Ay, Timothy L Bailey, and William Stafford Noble. Statistical confidence estimation for Hi-C data reveals regulatory chromatin contacts. Genome Research, 24(6):999–1011, 2014.

[3] Pablo G Cámara. Topological methods for genomics: present and future directions. Current Opinion in Systems Biology, 1:95–101, 2017.

[4] Pablo G Camara, Daniel IS Rosenbloom, Kevin J Emmett, Arnold J Levine, and Raul Rabadan. Topological data analysis generates high-resolution, genome-wide maps of human recombination. Cell Systems, 3(1):83–94, 2016.

[5] Gunnar Carlsson. Topological pattern recognition for point cloud data. Acta Numerica, 23:289–368, 2014.

[6] Mathieu Carriere and Raul Rabadan. Topological data analysis of single-cell Hi-C contact maps. 1812.01360, 2018.

[7] Giacomo Cavalli and Tom Misteli. Functional implications of genome topology. Nature Structural and Molecular Biology, 20(3):290–299, 2013.

[8] Joseph Minhow Chan, Gunnar Carlsson, and Raul Rabadan. Topology of viral evolution. Proceedings of the National Academy of Sciences, pages 18566–18571, 2013.

[9] Frédéric Chazal and Bertrand Michel. An introduction to topological data analysis: fundamental and practical aspects for data scientists. 1710.04019, 2017.

[10] Yu Chen, Yang Zhang, Yuchuan Wang, Liguo Zhang, Eva K Brinkman, Stephen A Adam, Robert Goldman, Bas van Steensel, Jian Ma, and Andrew S Belmont. Mapping 3D genome organization relative to nuclear compartments us-ing TSA-Seq as a cytological ruler. The Journal of Cell Biology, 217(11):4025–4048, 2018.

[11] Job Dekker, Karsten Rippe, Martijn Dekker, and Nancy Kleckner. Capturing chromosome confor-mation. Science, 295(5558):1306–1311, 2002.

[12] Jesse R Dixon, Inkyung Jung, Siddarth Selvaraj, Yin Shen, Jessica E Antosiewicz-Bourget, Ah Young Lee, Zhen Ye, Audrey Kim, Nisha Rajagopal, Wei Xie, et al. Chromatin architecture reorganization during stem cell differentiation. Nature, 518(7539):331–336, 2015.

[13] Geet Duggal, Rob Patro, Emre Sefer, Hao Wang, Darya Filippova, Samir Khuller, and Carl Kingsford. Resolving spatial inconsistencies in chromosome conformation measurements. Algorithms for Molecular Biology, 8:8, 2013.

[14] Geet Duggal, Hao Wang, and Carl Kingsford. Higher-order chromatin domains link eQTLs with the expression of far-away genes. Nucleic Acids Research, 42(1):87–96, 2014.

[15] Kevin Emmett, Benjamin Schweinhart, and Raul Rabadan. Multiscale topology of chromatin folding. 1511.01426, 2015.

[16] Paul A Fields, Vijay Ramani, Giancarlo Bonora, Galip Gurkan Yardimci, Alessandro Bertero, Hans Reinecke, Lil Pabon, William S Noble, Jay Shendure, and Charles Murry. Dynamic reorganization of nuclear architecture during human cardiogenesis. bioRxiv, page 222877, 2017.

[17] Mattia Forcato, Chiara Nicoletti, Koustav Pal, Carmen Maria Livi, Francesco Ferrari, and Silvio Bicciato. Comparison of computational methods for Hi-C data analysis. Nature Methods, 14(7):679–685, 2017.

[18] Timothy SC Hinks, Tom Brown, Laurie CK Lau, Hitasha Rupani, Clair Barber, Scott Elliott, Jon A Ward, Junya Ono, Shoichiro Ohta, Kenji Izuhara, et al. Multidimensional endotyping in patients with severe asthma reveals inflammatory heterogeneity in matrix metalloproteinases and chitinase 3–like protein 1. Journal of Allergy and Clinical Immunology, 138(1):61–75, 2016.

[19] Maxim Imakaev, Geoffrey Fudenberg, Rachel Patton McCord, Natalia Naumova, Anton Goloborodko, Bryan R Lajoie, Job Dekker, and Leonid A Mirny. Iterative correction of Hi-C data reveals hallmarks of chromosome organization. Nature Methods, 9(10):999–1003, 2012.

[20] Li Li, Wei-Yi Cheng, Benjamin S Glicksberg, Omri Gottesman, Ronald Tamler, Rong Chen, Erwin P Bottinger, and Joel T Dudley. Identification of type 2 diabetes subgroups through topological analysis of patient similarity. Science Translational Medicine, 7(311):311ra174, 2015.

[21] Erez Lieberman-Aiden, Nynke L van Berkum, Louise Williams, Maxim Imakaev, Tobias Ragoczy, Agnes Telling, Ido Amit, Bryan R Lajoie, Peter J Sabo, Michael O Dorschner, Richard Sandstrom, Bradley Bernstein, M A Bender, Mark Groudine, Andreas Gnirke, John Stamatoyannopoulos, Leonid A Mirny, Eric S Lander, and Job Dekker. Comprehensive mapping of long-range interactions reveals folding principles of the human genome. Science, 326(5950):289–293, 2009.

[22] Clément Maria, Jean-Daniel Boissonnat, Marc Glisse, and Mariette Yvinec. The Gudhi library: Simplicial complexes and persistent homology. In International Congress on Mathematical Software, pages 167–174. Springer, 2014.

[23] Monica Nicolau, Arnold J Levine, and Gunnar Carlsson. Topology based data analysis identifies a subgroup of breast cancers with a unique mutational profile and excellent survival. Proceedings of the National Academy of Sciences, pages 7265–7270, 2011.

[24] Benjamin D Pope, Tyrone Ryba, Vishnu Dileep, Feng Yue, Weisheng Wu, Olgert Denas, Daniel L Vera, Yanli Wang, R Scott Hansen, Theresa K Canfield, et al. Topologically associating domains are stable units of replication-timing regulation. Nature, 515(7527):402–405, 2014.

[25] Suhas S P Rao, Miriam H Huntley, Neva C Durand, Elena K Stamenova, Ivan D Bochkov, James T Robinson, Adrian L Sanborn, Ido Machol, Arina D Omer, Eric S Lander, and Erez Lieberman Aiden. A 3D map of the human genome at kilobase resolution reveals principles of chromatin looping. Cell, 159(7):1665–1880, Dec 2014.

[26] Sarah Rennie, Maria Dalby, Lucas van Duin, and Robin Andersson. Transcriptional decomposition reveals active chromatin architectures and cell specific regulatory interactions. Nature Communications, 9(1):487, 2018.

[27] Abbas H Rizvi, Pablo G Camara, Elena K Kandror, Thomas J Roberts, Ira Schieren, Tom Maniatis, and Raul Rabadan. Single-cell topological RNA-seq analysis reveals insights into cellular differentiation and development. Nature Biotechnology, 35(6):551, 2017.

[28] Nicolas Servant, Nelle Varoquaux, Bryan R Lajoie, Eric Viara, Chong-Jian Chen, Jean-Philippe Vert, Edith Heard, Job Dekker, and Emmanuel Barillot. HiC-Pro: an optimized and flexible pipeline for Hi-C data processing. Genome Biology, 16(1):259, 2015.

[29] Edwin H Spanier. Algebraic topology. 1966. MacGraw-Hill, New York, 1966.

[30] Larry Wasserman. Topological data analysis. Annual Review of Statistics and Its Application, 5:501–532, 2018.

[31] Y William Yu, Noah M Daniels, David Christian Danko, and Bonnie Berger. Entropy-scaling search of massive biological data. Cell systems, 1(2):130–140, 2015.

